# Bioorthogonal labeling with tetrazine-dyes for super-resolution microscopy

**DOI:** 10.1101/503821

**Authors:** Gerti Beliu, Andreas Kurz, Alexander Kuhlemann, Lisa Behringer-Pliess, Natalia Wolf, Jürgen Seibel, Zhen-Dan Shi, Martin Schnermann, Jonathan B. Grimm, Luke D. Lavis, Sören Doose, Markus Sauer

**Affiliations:** Department of Biotechnology and Biophysics, Biocenter, University of Würzburg, Am Hubland, 97074 Würzburg, Germany; Institute of Organic Chemistry, University of Würzburg, Am Hubland, 97074 Würzburg, Germany; Imaging Probe Development Center, National Heart, Lung, and Blood Institute, National Institutes of Health, Rockville, Maryland, 20850, USA; Chemical Biology Laboratory, Center for Cancer Research, National Cancer Institute, Frederick, MD 21702, USA; Janelia Research Campus, Howard Hughes Medical Institute, 19700 Helix Drive, Ashburn, Virginia 20147, USA

## Abstract

Genetic code expansion (GCE) technology allows the specific incorporation of functionalized noncanonical amino acids (ncAAs) into proteins. Here, we investigated the Diels-Alder reaction between trans-cyclooct-2-ene (TCO)-modified ncAAs, and 22 known and novel 1,2,4,5-tetrazine-dye conjugates spanning the entire visible wavelength range. A hallmark of this reaction is its fluorogenicity - the tetrazine moiety can elicit substantial quenching of the dye. We discovered that photoinduced electron transfer (PET) from the excited dye to tetrazine as the main quenching mechanism in red-absorbing oxazine and rhodamine derivatives. Upon reaction with dienophiles quenching interactions are reduced resulting in a considerable increase in fluorescence intensity. Efficient and specific labeling of all tetrazine-dyes investigated permits super-resolution microscopy with high signal-to-noise ratio even at the single-molecule level. The different cell permeability of tetrazine-dyes can be used advantageously for specific intra- and extracellular labeling of proteins and highly sensitive fluorescence imaging experiments in fixed and living cells.

---

Single-molecule localization microscopy is a powerful method for subdiffraction-resolution fluorescence imaging of cells and tissue^1,2^. Since the density of fluorophores controls the achievable structural resolution^3^, efficient and specific labeling with fluorescent probes is a decisive factor in this super-resolution microscopy technique. Despite recent progress in the development of new fluorophores with higher fluorescence quantum yield, photostability, and intrinsic photoswitching in aqueous buffer^4–8^, specific and efficient labeling of the molecule of interest remains a significant challenge. As the field moves toward ever higher spatial resolution using, e.g. expansion microscopy^9,10^, the effective size of the label (fluorophore, linker and affinity reagent) will be the main limiting factor of super-resolution microscopy.

Immunolabeling with antibodies is still the method of choice for fluorescence imaging of fixed cells, but the large size of antibodies introduces a displacement of 10-15 nm of the fluorophore from the molecule of interest^1^. This linkage error is smaller for fluorescent proteins and protein-based self-labeling tags but is still several nanometers^11^. Small (1.5 × 2.5 nm) camelid antibodies (‘nanobodies’) directed against green fluorescent protein (GFP) or shorter peptide epitopes have been used successfully for super-resolution microscopy^12^ but the palette of targets is small^13^. Even the most optimized labeling strategies using short peptide tags labeled with bivalent nanobodies or fluorescently labeled monomeric streptavidin yield linkage errors of ~ 2 nm in *direct* stochastic optical reconstruction microscopy (*d*STORM) experiments^14–17^. This problem demands the development of efficient labeling methods with small dyes, which can be site-specifically and quantitatively attached to a protein of interest with low linkage error.

Genetic code expansion (GCE) technology enables the introduction of noncanonical amino acids (ncAAs) with small functional groups at any position in a target protein^18–20^. In this strategy, a native codon is replaced with a rare codon, such as the amber (TAG) stop codon, at a specific site in the gene of the protein of interest. The modified protein is then expressed in mammalian cells along with an additional tRNA-tRNA synthetase pair (tRNA-RS) that is orthogonal to the host translational machinery. The active site of the tRNA synthetase enzyme is engineered to only accept a specific ncAA, which is incorporated into a tRNA that recognizes the rare codon. The ncAA is simply added to the growth medium and thereby incorporated into the protein at a specific site^18–20^. A particularly promising type of ncAA include strained alkenes, such as trans-cyclooct-2-ene (TCO*), that can react with a 1,2,4,5-tetrazine in an ultrafast, specific, and bioorthogonal inverse electron-demand Diels-Alder reaction. In this strategy, TCO*-modified ncAAs, such as TCO*-L-lysine (TCO*-Lys), can be efficiently and directly labeled with organic dyes with minimal linkage error^21^. The high selectivity and rate of this click chemistry reaction has resulted in a large number of commercially available fluorophore-tetrazine conjugates allowing labeling of mammalian cells and whole organisms with organic dyes, even in living systems^22–24^.

Another interesting property of this labeling strategy is the potential for fluorogenicity. It has been reported that some tetrazine-functionalized dyes (tetrazine-dyes) function as fluorogenic probes, meaning these compounds substantially increase fluorescence intensity upon reaction with the strained dienophile such as TCO^25,26^. This makes tetrazine-dyes especially interesting for live-cell labeling and fluorescence imaging applications since the fluorogenic reaction could lower background and potentially eliminate the need for washing out excess fluorophore. Despite these apparent advantages however, the use of ncAAs and tetrazine-dyes to label proteins for super-resolution microscopy applications remain rare and unoptimized^27–29^.

Motivated by these considerations, we studied the spectroscopic characteristics and quenching mechanism of 22 known and novel tetrazine-dyes that span the entire visible spectral range. The tetrazine-dyes included in this study are ATTO425, ATTO465, ATTO488, ATTO532, Cy3, carboxytetramethylrhodamine (TAMRA), ATTO550, ATTO565, ATTO590, ATTO594, ATTO620, Si-rhodamine (SiR)^4^, ATTO647N, Cy5, ATTO655, ATTO680, ATTO700, two large Stokes shift dyes AZ503 and AZ519, as well as the recently introduced Si-rhodamine derivative JF_646_^6^, spontaneously blinking HMSiR^5^, and the bridged carbocyanine dye Cy5B^8^ (Supplementary Fig. S1).

## Results and discussion

### Spectroscopic characteristics of tetrazine-dyes

The absorption and emission maxima of the 22 tetrazine-dyes were measured in aqueous buffer (PBS, pH 7.4) before and after reaction with ncAA TCO*-Lys (Table 1). In TCO*-Lys isomerization to the less reactive *cis*-isomer is prevented by stabilizing the compound for several days under physiological conditions^30^. The majority of tetrazine-dyes are commercially available as 3-methyl-6-phenyl-1,2,4,5-tetrazine (Me-Tet) derivatives and only some of them are also available as 3-phenyl-1,2,4,5-tetrazine (H-Tet) derivatives. Those dyes which are not available as tetrazine derivatives were synthesized by reacting W-hydroxysuccinimide derivatives of the fluorophores with 3-(*p*-benzylamino)-1,2,4,5-tetrazine (H-Tet-amine; Supplementary Fig. S2). The main difference between Me-Tet and H-Tet dye conjugates are the different reactivity and chemical stability: H-Tet exhibits an approximately 30-fold higher click reaction rate constant but a lower chemical stability^30^.

**Table 1.**
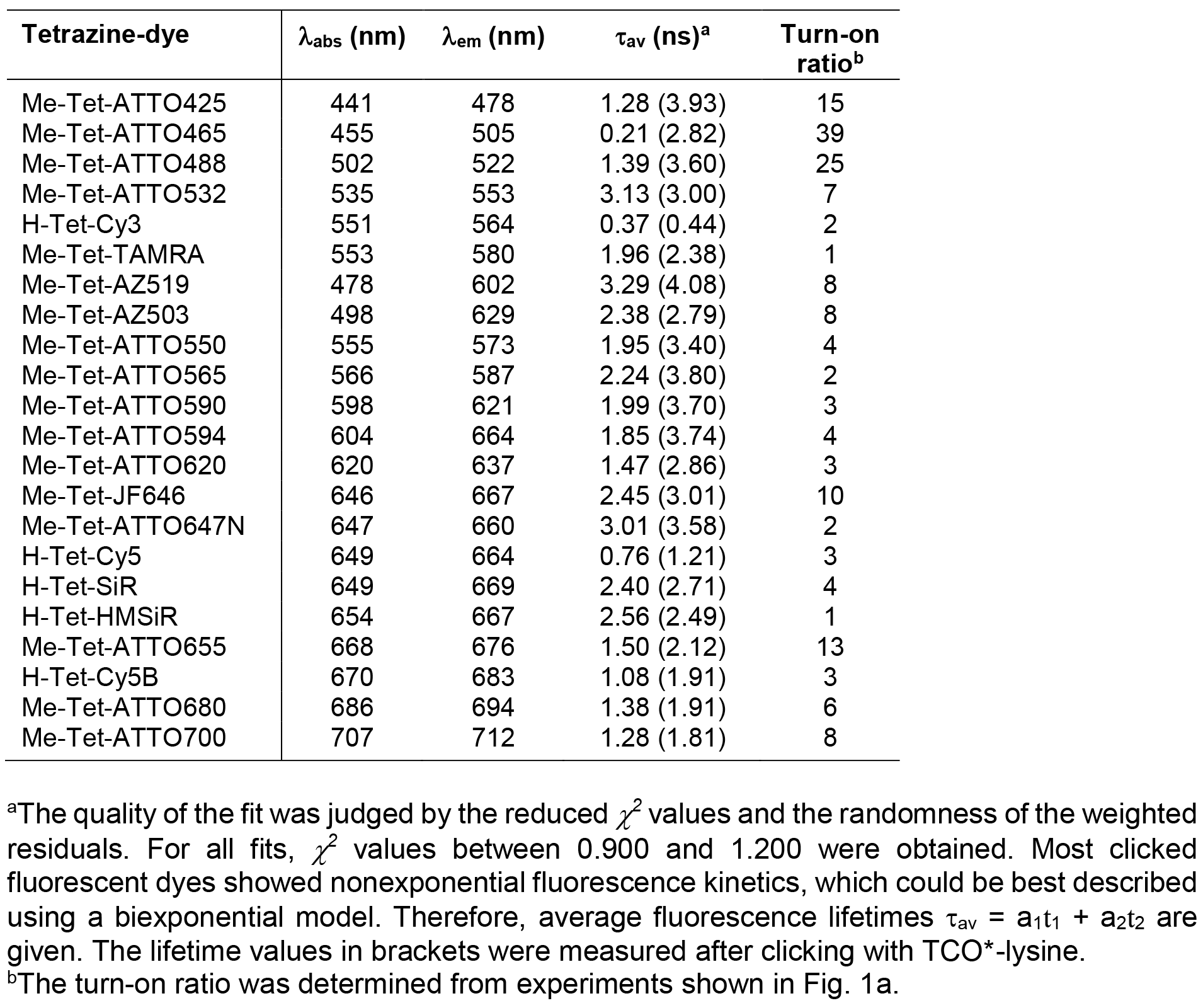
Spectroscopic characteristics (absorption maxima, λ_abs_ (nm), emission maxima, λ_em_ (nm), fluorescence lifetimes, τ (ns)) of flexibly linked tetrazine-dyes after click reaction with TCO*-lysine and turn-on factors measured in PBS, pH 7.4. Click reactions were performed by adding 25 μM TCO*-Lys to 1 μM dye solutions.

Almost all tetrazine-dyes investigated exhibited a significant fluorescence increase upon reaction with TCO*-Lys (Table 1, Supplementary Fig. S3). The increase in fluorescence intensity is most pronounced for the shorter wavelength absorbing dyes ATTO425, ATTO465, and ATTO488 with turn-on ratios of 15-40. Due to the broad absorption spectrum of tetrazine peaking at ~ 515 nm it can efficiently quench the fluorescence of dyes emitting at wavelengths ≤ 550 nm by fluorescence resonance energy transfer (FRET) (Supplementary Fig. S2)^25^. Hence, the tetrazine chromophore can act as both quencher and bioorthogonal click-reaction group. Indeed, the turn-on ratios of the tetrazine-dyes ATTO425, ATTO465, ATTO488, and ATTO532 after coupling to TCO*-lysine scale according to the emission maxima of the dyes and their overlap with the absorption spectrum of tetrazine (Table 1, Supplementary Figs. S1,S2).

The longer wavelength absorbing oxazine dyes ATTO655, ATTO680, and ATTO700, and the large Stoke-shift dyes AZ503 and AZ519 show turn-on ratios of 6-13 (Table 1, Fig. 1a, Supplementary Fig. S3). These high turn-on ratios are not entirely reflected in the increase in fluorescence lifetime indicating that short fluorescence lifetime components are missed in our time-correlated single-photon counting (TCSPC)-experiments with a time resolution of approximately 40 ps.

**Figure 1.**
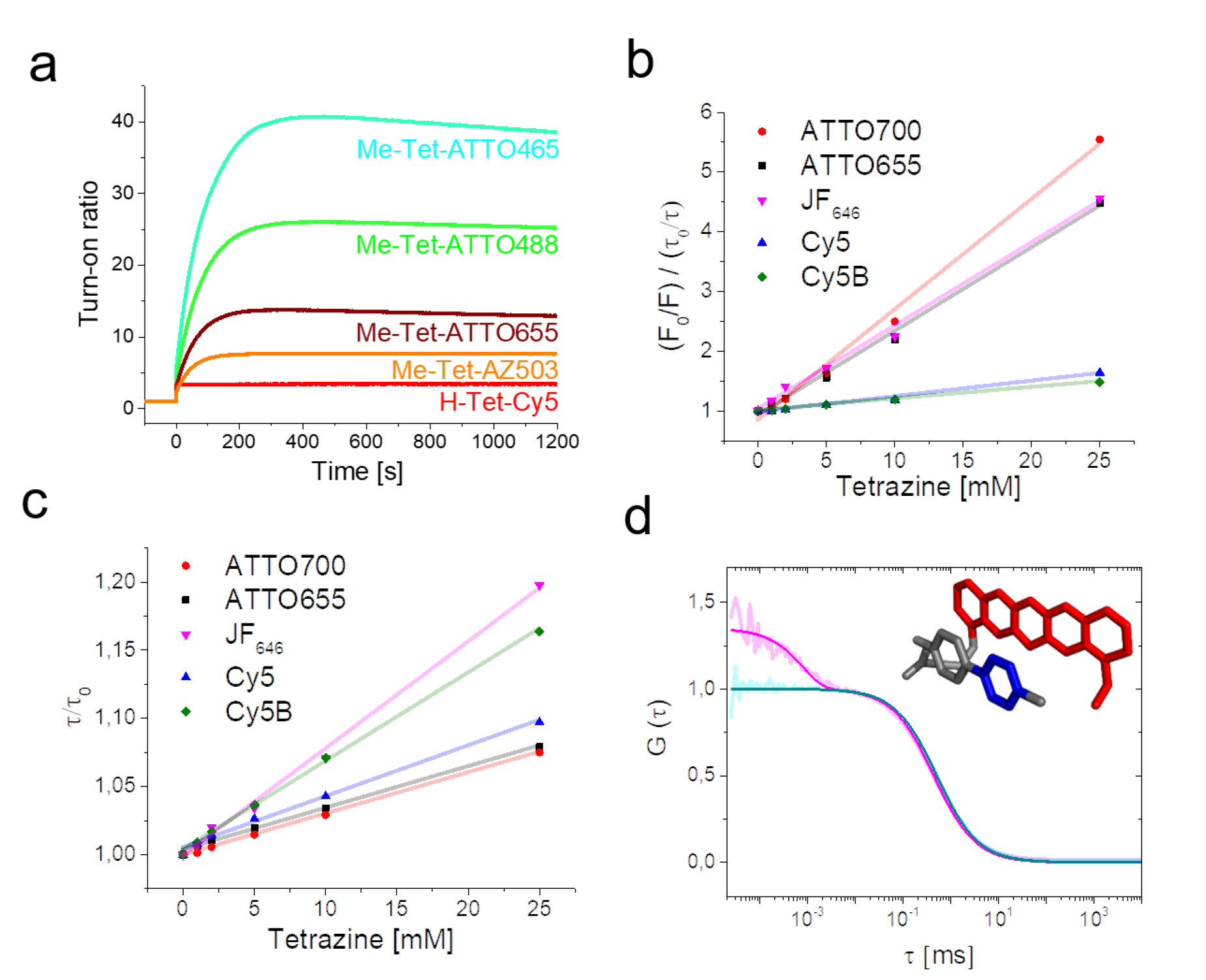
Fluorescence quenching of flexibly-linked tetrazine dyes. **a**. Fluorescence turnon of 1 μM solutions of different tetrazine-dyes upon addition of 25 μM TCO*-Lys. **b**. Static and **c**. dynamic quenching of ATTO655, ATTO700, Cy5, Cy5B, and JF_646_ with 0 - 25 mM Me-Tet-amine. **d**. FCS-curve of Me-Tet-ATTO655 recorded in PBS, pH 7.4 in the absence and presence of 10 μM TCO*-Lys. All measurements were performed in PBS, pH 7.4 at room temperature (25°C). Inset: Model for formation of weak or nonfluorescent ground-state complexes between an oxazine dye and tetrazine. Due to the flexible linker the oxazine dye (e.g. ATTO655) (red) can adopt a nearly coplanar stacking conformation with respect to tetrazine (blue).

Long-term spectroscopy studies revealed that some dyes such as ATTO465, ATTO488, and ATTO655 show a slight but highly reproducible decrease in fluorescence intensity after the initial strong increase upon addition of TCO*-Lys (Fig. 1a, Supplementary Fig. S3), which can be attributed most likely to tautomerization of the TCO*-tetrazine conjugate resulting in the release of lysine by decarboxylative elimination^31^.

Comparative click reactions demonstrate that the click reaction of H-Tet-ATTO488 and H-Tet-Cy5 with TCO*-Lys proceeds approximately 30-times faster than for the corresponding Me-Tet derivatives^32^ but the turn-on ratios differ significantly especially for the two ATTO488 tetrazine derivatives (Supplementary Fig. S4). In particular, H-Tet-ATTO488 is substantially less quenched than Me-Tet-ATTO488. Accordingly, the turn-on ratio is much higher for the Me-Tet derivative. Assuming similar FRET efficiencies for both ATTO488 conjugates the different turnon ratios can only be explained by the reduced chemical stability of H-Tet^32^. Obviously, a fraction of H-Tet-ATTO488 decomposes spontaneously in physiological buffer because of the higher reactivity, which results correspondingly in a lower FRET-efficiency and turn-on ratio (Supplementary Fig. S4).

Higher quenching efficiencies with turn-on rates of > 1,000 have been achieved in the green wavelength region by using through-bond energy transfer (TBET) in tetrazine-BODIPY derivatives^26^. However, efficient TBET quenching has been restricted to tetrazine-dyes absorbing in the blue-green wavelength range, e.g. bodipy and coumarin dyes, which are less suited for super-resolution microcopy^25,26,33^. Flexibly-linked tetrazine-dyes as used here are easier to access from a synthetic standpoint but exhibit lower turn-on ratios after reaction with dienophiles (Table 1)^27^.

### Fluorescence quenching in red-absorbing tetrazine-dyes

To shed light on the fluorescence quenching mechanism in the longer wavelength absorbing flexibly-linked tetrazine-dyes we performed intermolecular steady-state and time-resolved fluorescence quenching experiments with free dyes and Me-Tet-amine under physiological conditions (PBS, pH 7.4) following previous studies on tryptophan quenching^34,35^ (Figs. 1b, 1c, Supplementary Fig. S5). The time-resolved fluorescence decays remain almost monoexponential with a longer fluorescence lifetime, dependent on the quencher concentration, and an additional shorter decay component with lifetimes in the range of a few tens of picoseconds. For the quencher dependent longer fluorescence lifetimes, we observed linear Stern-Volmer plots for all measured dyes (ATTO655, ATTO700, JF_646_, CY5, and Cy5B) (Fig. 1c) and calculated bimolecular dynamic quenching rate constants *k_dyn_* of (1.7±0.1) × 10^9^ M^-1^S^-1^, (1.9±0.1) × 10^9^ M^-1^S^-1^, (2.6±0.1) × 10^9^ M^-1^S^-1^, (4.1±0.2) × 10^9^ s^-1^, and (3.5±0.1) × 10^9^ M^1^s^-1^, respectively. These values represent dynamic quenching that is approaching the diffusion limit.

In addition, strong static quenching was observed for the two oxazine dyes ATTO655 and ATT0700 as well as for the Si-rhodamine dye JF_646_ (Fig. 1b) suggesting that non- or only weakly fluorescent ground-state complexes between the longer wavelength absorbing oxazine and Si-rhodamine dyes and Me-Tet-amine form in aqueous solutions, where the lifetime of the complexes is below the time-resolution of our TCSPC setup.

From the static quenching component the association constant, *K_a_*, for complex formation can be calculated by plotting (F_0_/F)/(τ_0_/τ) versus the quencher (Me-Tet-amine) concentration (Fig. 1b, Supplementary Fig. S5). We estimated *K_a_* for complex formation between the dyes ATTO655, ATT0700, JF_646_ and Me-Tet-amine to be (135±4) M^-1^, (175±7) M^-1^, and (140±3) M^-1^, respectively. Cy5 and Cy5B, on the other hand, show only weak static quenching (Supplementary Fig. S5) with *K_a_* of (24±1) M^-1^, and (19±1) M^-1^, respectively.

Me-Tet-ATTO655, Me-Tet-ATTO680, Me-Tet-ATTO700, and Me-Tet-JF_646_ show a high turnon ratio (Table 1) and are therefore efficiently quenched by Me-Tet-amine also in intramolecular quenching experiments. This finding indicates that the dye and the tetrazine moiety, when connected by a flexible linker, can adopt a nearly coplanar stacking conformation, which is required for efficient fluorescence quenching (Fig. 1d).

Our results indicate that the underlying quenching mechanism in longer wavelength absorbing oxazine and Si-rhodamine dyes is only efficient at very short distances, i.e. within complexes. This is also supported by the observation that the fluorescence intensity of ATTO655, ATTO680, and ATTO700 tetrazine conjugates increases strongly upon addition of denaturing agents such as guanidinium chloride while ATTO488 and Cy5 show only small effects if at all (Supplementary Fig. S6). Hence, it can be concluded that hydrophobic interactions between the longer wavelength absorbing oxazine and Si-rhodamine dyes and Me-Tet with a stacked arrangement of the conjugated π-electron systems play an important role in the formation of ground-state complexes with ultrafast fluorescence quenching. Since tetrazine exhibits a very high electron affinity^36,37^, the excited longer-wavelength absorbing dyes are most likely efficiently quenched in their stacked conformation via photoinduced electron transfer (PET). The carbocyanine dyes such as Cy5 exhibit a higher water solubility and are thus less prone to form complexes with Me-Tet. In the case of Me-Tet-ATTO488 fluorescence quenching is dominated by through-space FRET where complex formation between dye and tetrazine moiety does not play a dominant role.

### Investigating fluorescence quenching dynamics of tetrazine-dyes by fluorescence correlation spectroscopy (FCS)

Knowing that ATTO655 and similar dyes form nonfluorescent complexes with tetrazine in aqueous solutions driven by hydrophobic interactions, we can assume an equilibrium between two states for the dye in conformationally flexible tetrazine-dyes: an ‘open’ fluorescent state *A_open_* and a ‘closed’, complexed nonfluorescent state *B_closed_*. Both states are populated according to the closing and opening rate constants, *k_closing_* and *k_opening_*. To investigate this equilibrium in more detail, we performed fluorescence correlation spectroscopy (FCS) experiments (Fig. 1d). FCS analyzes temporal fluorescence fluctuations from highly diluted samples (nanomolar concentrations), probing molecules diffusing through a confined detection volume (typically 1 femtoliter with 1-20 molecules) by Brownian motion^38^. Characteristic time scales of molecular processes that result in fluctuating fluorescence emission can be measured under thermodynamic equilibrium conditions with nanosecond time resolution^39–41^.

Under moderate excitation conditions, no photophysical process, such as intersystem crossing, is apparent in the submillisecond time domain of FCS-curves recorded from Me-Tet-ATTO655 after reaction with TCO*-Lys (Fig. 1d). The FCS curve shows a millisecond decay corresponding to the molecules’ diffusion through the confocal excitation/detection volume. However, the FCS curve recorded from Me-Tet-ATTO655 before addition of TCO*-Lys displays an additional fast decay occurring on the nano-to microsecond time scale together with a change in amplitude (Fig. 1d). Bimolecular experiments with ATTO655 and 25 mM tetrazine reveal a similar FCS decay (Supplementary Fig. S7) confirming the idea of diffusion-driven formation of non-fluorescent complexes. By fitting an analytical FCS curve representing two-state fluorescence intermittency to the bimolecular data with a mixture of free ATTO655 and tetrazine we estimated *k_closing_* = (0.04±0.01) × 10^9^ s^-1^ and *k_opening_* = (0.015±0.002) × 10^9^ s^-1.^ These values represent an association rate constant of (1.5±0.5) × 10^9^ M^-1^s^-1^ and an association constant of (100±23) M^-1^, close to the values from Stern-Volmer analysis. Concentration estimates from the amplitude of FCS curves confirm that all fluorophores contribute to the observed bimolecular FCS decays and no population of fluorophores exist, that are quenched over time periods longer than ~1 ms (the diffusion-limited observation time) (Supplementary Fig. S7).

It is thus reasonable, that the nano-to microsecond decay recorded for Me-Tet-ATTO655 reflects fluorescence fluctuations caused by intramolecular quenching of ATTO655 upon contact with Me-Tet mediated by the conformational flexibility of the linker. The time scale of the fast quenching fluctuations is well separated from that of translational diffusion. The calculated rate constants for opening and closing, *k_opening_* of ~ 9,0 × 10^6^ s^-1^ and *k_closing_* of ~ 3,3 × 10^6^ s^-1^, demonstrate that ATTO655 stays on average 27% of the time in its quenched complexed conformation. The fast decay thus accounts for about one fourth of the observed turn-on ratio of 13 (Table 1). The remaining turn-on ratio is indeed reflected in a concentration difference by about a factor of four that is observed from the amplitude of FCS curves (Supplementary Fig. S7). This observation of multiple decay components on time scales above and below the FCS diffusion time of ~1 ms indicates that the intramolecular linker introduces additional conformational constraints for the process of complex formation.

Whereas Me-Tet-ATTO680 and Me-Tet-ATTO700 show similar behavior, the FCS-curves recorded from Me-Tet-Cy5 and Me-Tet-Cy5B in the absence and presence of TCO*-Lys are identical demonstrating that quenched complexes are not formed. The observed fast decay for Cy5 is due to well-characterized cis/trans-isomerization (Supplementary Fig. S8)^42^. Accordingly, we observed that Cy5B, which is not capable of isomerization due to the constrained conjugated bond structure, does not show the isomerization decay (Supplementary Fig. S8).

### Super-resolution microscopy with tetrazine-dyes

In order to enable testing of different tetrazine-dyes for super-resolution microscopy applications in an experimentally easy and comparable way, we synthesized phalloidin-TCO^43^, for labeling of the actin skeleton of fixed cells and subsequent attachment of tetrazine-dyes by click chemistry. Confocal fluorescence images demonstrate that all tetrazine-dyes are well-suited for high-end microscopy (Fig. 2, Supplementary Fig. S9).

**Figure 2.**
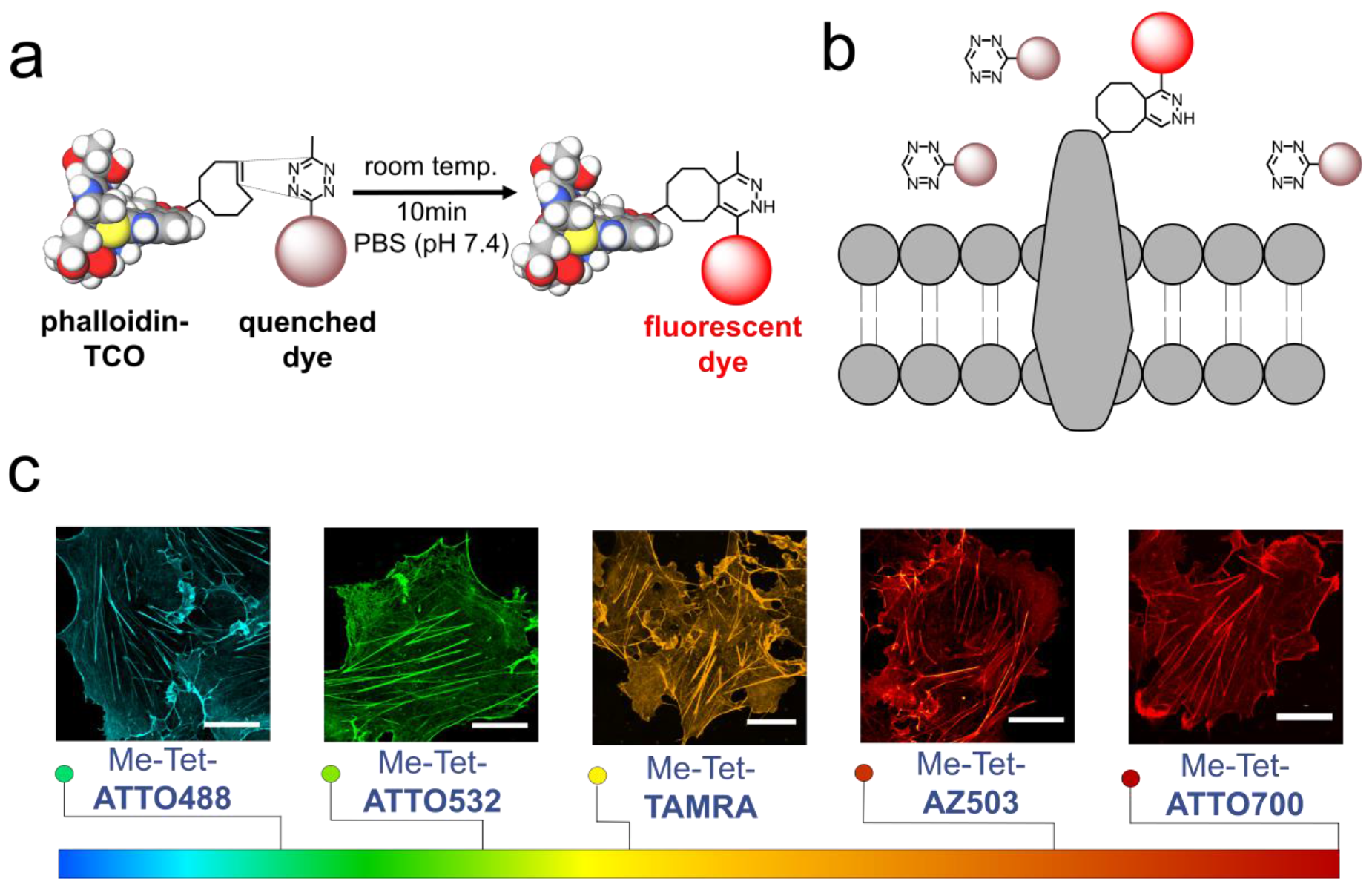
High-end microscopy of actin with tetrazine-dyes. **a**. Cos-7 and NIH-3T3 cells were fixed and labeled with phalloidin-TCO followed by click-labeling with different tetrazine-dyes. All tetrazine-dyes investigated show specific labeling of the actin skeleton. **b**. Tetrazine-dyes can also be used for site-specific labeling of the extracellular domain of membrane receptors as shown in Fig. 5. **c**. Various tetrazine-dyes spanning the entire visible part of the electromagnetic spectrum can be used successfully for confocal laser scanning microscopy of intracellular actin structures (see also Fig. 3). Scale bars, 20 μm.

Next, we tested their performance in super-resolution microscopy by re-scan confocal microscopy (RCM)^44,45^ and structured illumination microscopy (SIM)^46^ providing a resolution improvement factor of 1.4 and 2.0, respectively (Fig. 3, Supplementary Figs. S10 and S11). All tetrazine-dyes investigated, including the large Stokes-shift dyes AZ519 and AZ503 as well as the new dyes HMSiR^5^, JF _646_^6,7^, and Cy5B^8^, show excellent labeling efficiency and provide super-resolved actin images with high signal-to-noise ratio. Furthermore, fluorogenic tetrazine-dyes such as Me-Tet-ATTO488, Me-Tet-ATTO655, and Me-Tet-ATTO680 with turn-on ratios ≥ 6 (Table 1) enable wash-free super-resolution fluorescence imaging (Fig. 3b)^28,29^. Furthermore, also other tetrazine-dyes such as H-Tet-Cy3 and H-Tet-Cy5 with lower turn-on ratio allow wash-free fluorescence imaging (Supplementary Fig. S12). This finding demonstrates that not only the turn-on ratio but likewise a high water-solubility and low tendency of unspecific binding to cellular components in combination with superior click reactivity can promote wash-free imaging of intracellular structures.

Next, we performed single-molecule localization microscopy experiments to verify that tetrazine-dyes can be used as well for single-molecule sensitive fluorescence imaging (Fig. 4). The obtained super-resolution images demonstrate, that H-Tet-Cy5 coupled to phalloidin-TCO can be reliably photoswitched in the presence of millimolar concentrations of thiols enabling *d*STORM imaging^14,15^ of the actin skeleton with superior spatial resolution (Fig. 4a). In addition, we tested also the bridged carbocyanine dye Cy5B^8^ for PALM-imaging under reductive imaging conditions (Fig. 4b) and the spontaneously blinking dye HMSiR^5^ in PBS, pH 7.4 without addition of photoswitching buffer (Fig. 4c) and achieved similar image qualities; or to be precise, the dyes exhibit slightly different localization precisions of ~12 nm (Cy5), ~13 nm (HMSiR), and ~11 nm (Cy5B) due to the different localization intensities of 1 : 0.42 : 1.8 (Cy5 : HMSiR : Cy5B) recorded under the different experimental conditions.

**Figure 3.**
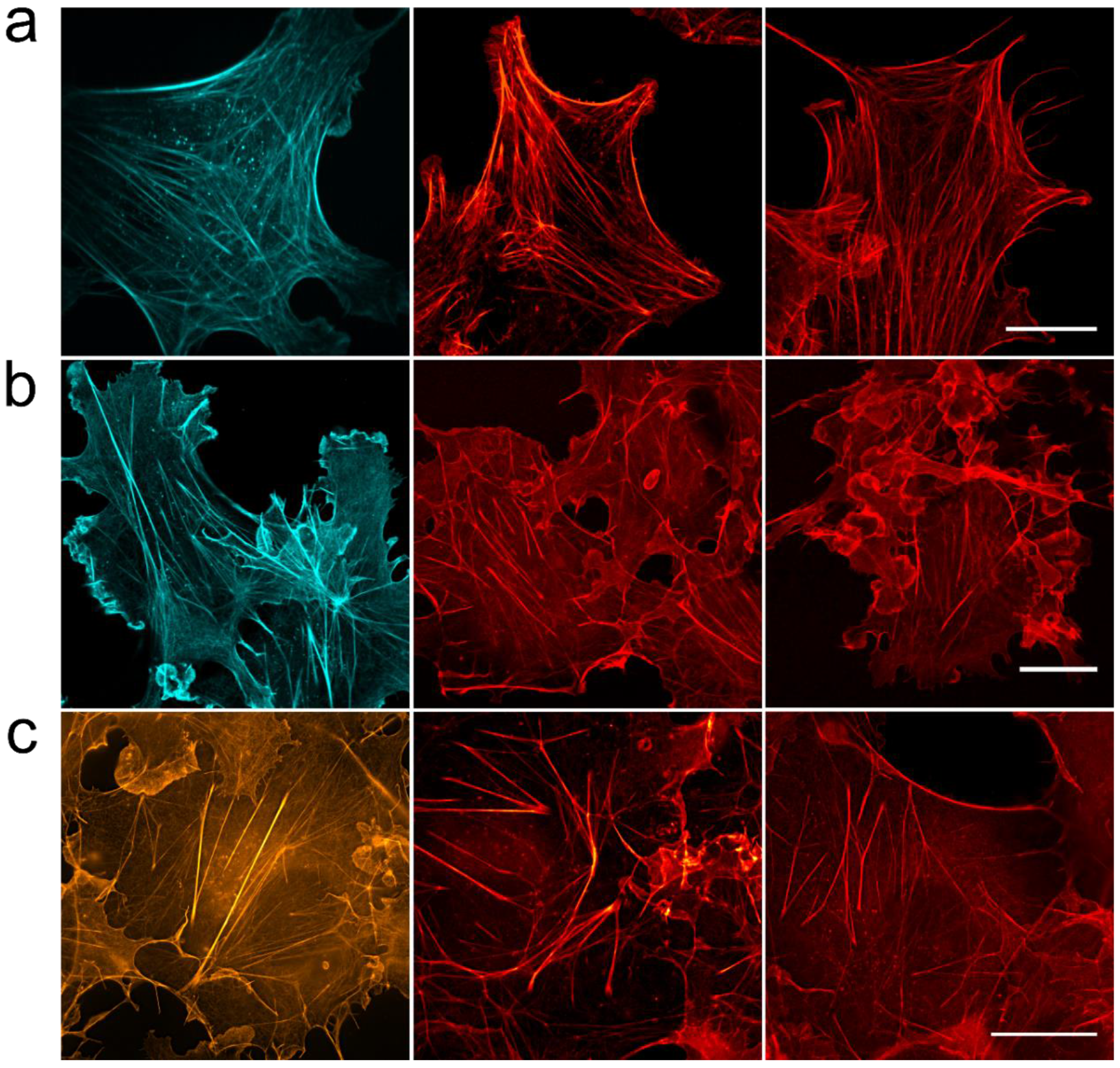
Super-resolution microscopy with tetrazine-dyes. a. RCM-images of actin in fixed NIH-3T3 cells labeled with phalloidin-TCO and Me-Tet-ATTO425, large Stokes-shift dyes Me-Tet-AZ503, and Me-Tet-SiR are shown from left to right. **b**. Wash-free confocal laser scanning microscopy images of fixed Cos-7 cells labeled with Me-Tet-ATTO488, Me-Tet-ATTO655, and Me-Tet-ATTO680 (from left to right). Cells were fixed, 1 h incubated with phalloidin-TCO, rinsed with PBS and then labeled with 3 μM tetrazine-dye and imaged directly after 10 min. without any washing step in the presence of unreacted free tetrazine-dye. **c**. SIM-images of actin in fixed Cos-7 cells labeled with phalloidin-TCO and Me-Tet-ATTO565, Me-Tet-ATTO620, and Me-Tet-JF_646_ are shown from left to right. Scale bars, 20 μm.

**Figure 4.**
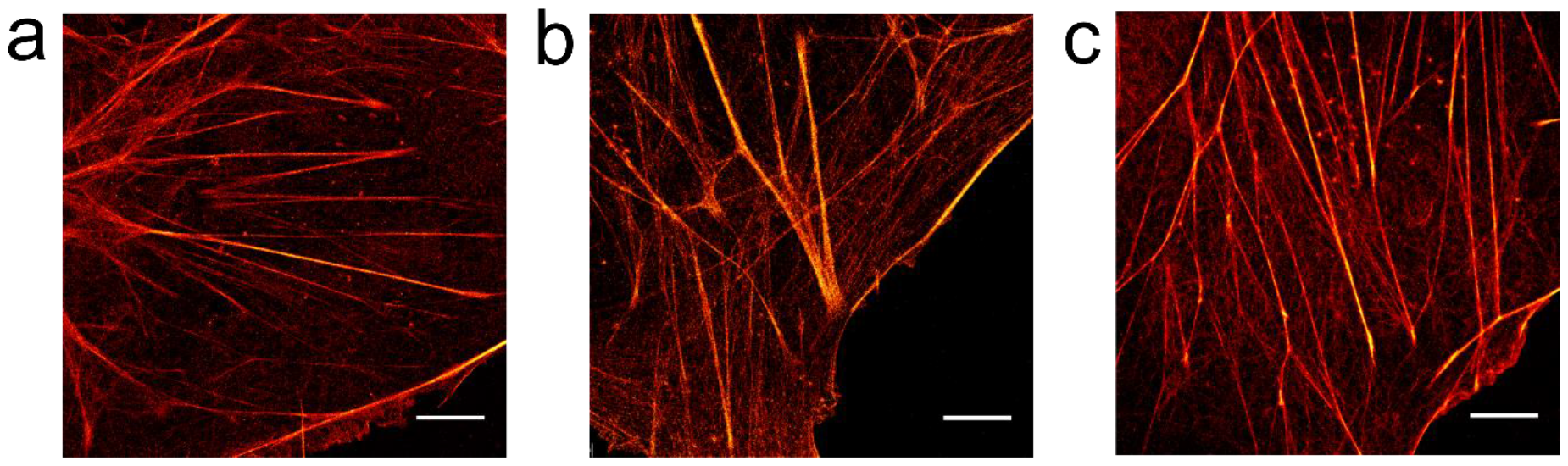
Single-molecule localization microscopy with tetrazine-dyes. Cos-7 cells were labeled with phalloidin-TCO and different tetrazine-dyes. **a**. *d*STORM-image of actin labeled with H-Tet-Cy5 in the presence of photoswitching buffer. **b**. PALM-image^8^ of actin labeled with H-Tet-Cy5B in PBS, pH 7.4. Cy5B was reduced with 0.1% NaBH4 for 10 min before imaging. **c**. Actin was labeled with phalloidin-TCO and H-Tet-HMSiR, a spontaneously blinking silicon rhodamine dye^5^, and imaged in PBS, pH 7.4. Irradiation was performed in all three examples using solely 640 nm laser light at irradiation intensities of ~ 2 kW/cm^2^. Scale bars, 5 μm.

### Visualization of membrane receptors by bioorthogonal labeling with tetrazine-dyes

To demonstrate the usefulness of click chemistry and tetrazine-dyes for fluorescence imaging of cells, we used GCE for site-specific labeling of extracellular domains of two membrane receptors: the kainate receptor (GluK2) and the tumor necrosis factor receptor 1 (TNFR1). Kainate receptors are ionotropic glutamatergic receptors that mediate fast excitatory neurotransmission and are localized to the presynaptic and postsynaptic sides of excitatory synapses^47^. The TNFR superfamily is a very important class of signal transduction molecules in the immune system^48^. TNFRs are single-membrane-spanning proteins that contain an extracellular TNF-binding region and a cytoplasmic tail.

To improve the incorporation efficiency of ncAAs and reduce the background signal we used a recently optimized pyrrolysine-based CGE system for click chemistry^49^. By adding a strong nuclear export signal (NES) to the N-terminus of the PylRS^AF^ sequence (NESPylRS^AF^), cytoplasmic localization of the protein of interest is promoted^49^. In combination with cell-impermeable dyes such as H-Tet-Cy5 this approach enables selective and efficient labeling of extracellular domains of correctly incorporated membrane receptors.

First, we exchanged serine at position 47 by TCO*-Lys in the extracellular domain of TNFR1 (TNFR1-tdEOS^S47TAG^) in HEK293T cells by amber suppression. We tested different H-Tet-Cy5 concentrations ranging from 10 nM to 5 μM and found that labeling with 1.5 μM H-Tet-Cy5 for 10 minutes is optimal. Cells were labeled with H-Tet-Cy5 and imaged by confocal laser scanning microscopy. With tdEOS attached C-terminally after the amber suppression site, the fluorescence signal of the fluorescent protein is only detectable when the ncAA is incorporated properly into the protein of interest. Live-cell fluorescence images clearly show selective labeling of TNFRs in the plasma membrane of cells (Figure 5a). Next, we labeled the kainate receptor GluK2 (GluK2^S343TAG^-eGFP) in HEK293T cells by GCE (amber suppression) with TCO*-Lys and H-Tet-Cy5. Live-cell confocal fluorescence images show again exclusively extracellular fluorescence labeling of GluK2 (Figure 5b). Corresponding đSTORM images demonstrate that GluK2 is expressed at high expression rates but homogeneously distributed in the plasma membrane of fixed HEK293T cells (Figure 5c).

**Figure 5.**
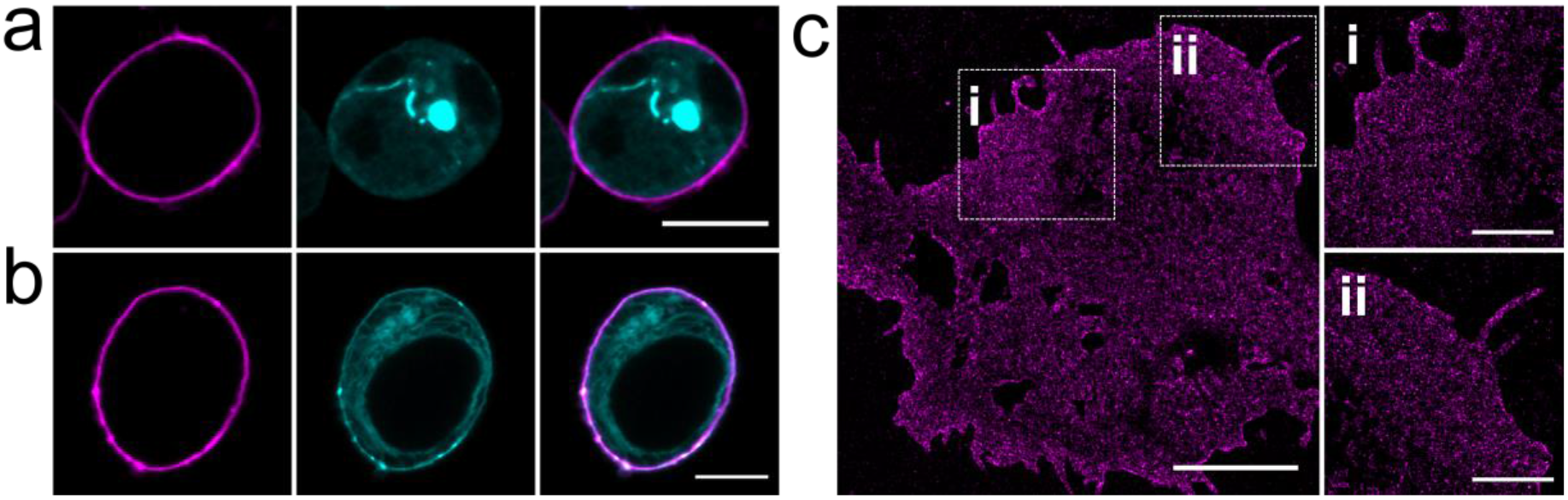
Visualization of membrane receptors by GCE and bioorthogonal labeling with tetrazine-dyes. **a**. Confocal fluorescence image of the clickable TNF receptor 1 (TNFR1^S47TAG^-tdEOS) and **b**. Kainate receptor (GluK2^S343TAG^-peGFP) in HEK293T cells. Site-specific extracellular labeling was performed with the cell impermeable H-Tet-Cy5 (1.5 uM) (magenta). The tdEOS and GFP channels, respectively, are shown in cyan. **c**. *d*STORM image of membrane receptors GluK2^S343TAG^ expressed in HEK293T cells shown a homogeneous distribution of receptors in the plasma membrane. Scale bars, 10 μm (a,b,c), 2 μm (expanded images i and ii).

### Live-cell bioorthogonal labeling

So far intracellular live-cell bioorthogonal labeling by GCE and fluorescence imaging has been hampered by the high fluorescence background resulting from the excess of unincorporated ncAAs^30^. Even water-soluble tetrazine-dyes such as H-Tet-SiR^4,30^ or tetrazine-dyes with high turn-on ratio do not automatically permit high-end live-cell fluorescence imaging of intracellular target proteins. In general, incorporation of TCO*-Lys into proteins in response to an in-frame amber stop codon does not alter cell viability. Depending on the incorporation site the expression level and cellular location might, however, be influenced. This off-target binding further increases the fluorescence background.

To demonstrate live-cell bioorthogonal labeling and imaging with tetrazine-dyes we selected tubulin as an important cytoskeleton component of cells in our experiments. Microtubules are dynamic structures composed of αβ-tubulin molecules that are constantly integrated or degraded as the microtubules grow and shorten. Microtubule dynamics can be easily monitored in live cells using fluorescently labeled tubulin and video microscopy. Since overexpression of α-tubulin can potentially influence the dynamics of endogenous microtubules after incorporation and generate a high fluorescent background^28^, we used two alternative live-cell tubulin labeling strategies. First, we introduced TCO*-Lys into the microtubule-associated protein (MAP) enscosine (E-MAP-115) which is known to be better tolerated by cells at high expression levels^50^ and second, we synthesized Docetaxel-TCO, a microtubule-stabilizing antimitotic drug^51–53^ (Fig. 6a). It has been shown that cells expressing four to ten times the physiological level of endogenous MAP exhibited microtubule dynamics indistinguishable from those of untransfected cells indicating that E-MAP-115 most likely serves to modulate microtubule functions or interactions with other cytoskeletal elements^50^. We inserted TCO*-Lys into the microtubule binding domain (EMTB) of E-MAP-115, which is C-terminally tagged with three GFPs (EMTB^87TAG^-3xGFP)^54^ for live-cell bioorthogonal labeling and fluorescence imaging (Fig. 6a). Thus, monitoring of the GFP signal allowed us to select strongly expressing cells and change to TCO*-Lys free medium one day before bioorthogonal labeling with cell-permeable H-Tet-SiR^4,28^ in order to minimize the non-specific background signal. In addition, we used again the tRNAPyl/NESPylRS^AF^ pair to improve nuclear export of tRNA-Synthetase^49^.

**Figure 6.**
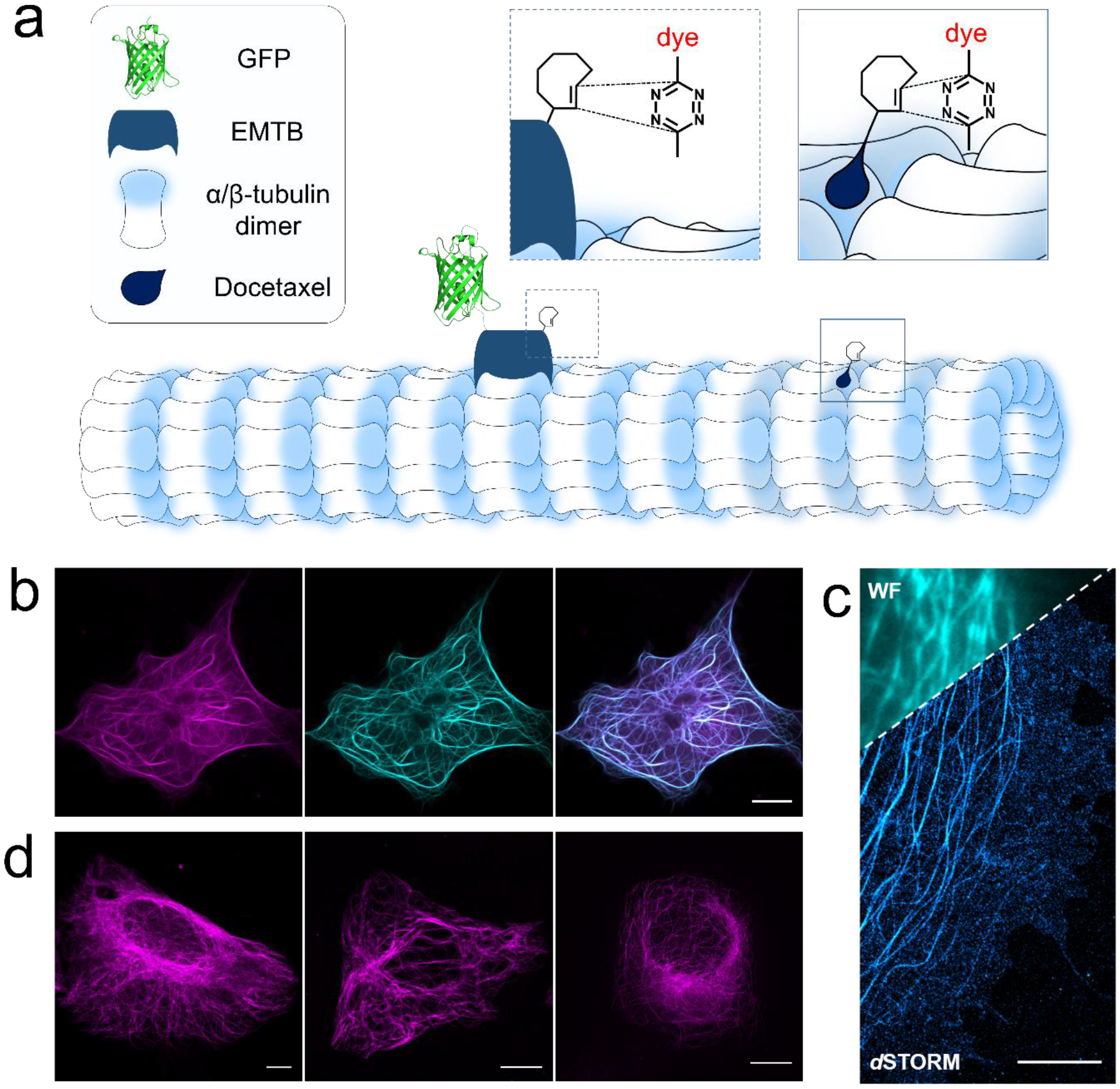
Intracelullar live-cell labeling and imaging of clickable microtubule-associated protein (EMTB) in COS-7 cells. a. Scheme of incorporation of unnatural amino acid TCO*-Lys into the microtubule-associated protein EMTB^87TAG^-3xGFP via GCE and bioorthogonal labeling with a tetrazine-dye. Alternatively, microtubules can be labeled in living cells with Docetaxel-TCO followed by click labeling with a cell permeable tetrazine-dye. **b**. Live-cell confocal fluorescence images of the construct EMTB^87TAG^-3xGFP labeled with 3 μM of the membrane-permeable H-Tet-SiR for 10 min (GFP: green, SiR: magenta, and overlay). **c**. Single-molecule localization microscopy image of the same construct. Cells were fixed and labeled with 3 μM H-Tet-HMSiR and then imaged in PBS (pH 7.4). The upper left corner shows an overlay with the corresponding widefield image (WF). **d**. Live cell confocal, re-scan confocal, and SIM fluorescence images (from left to right) of U2OS cells treated with 10 μM Docetaxel-TCO for 30 min and labeled with 10 μM H-Tet SiR for 10 min. Scale bars, 5 μm.

Using these optimized approach, we were able to perform fluorescence imaging of microtubule dynamics in living cells by re-scan confocal microscopy (RCM)^44,45^ (Fig. 6b, Supplementary Videos 1 and 2). However, additional single-molecule localization microscopy experiments of fixed EMTB^87TAG^-3xGFP cells with H-Tet-HMSiR illustrate that the fluorescence background of unbound EMTB still limits the achievable image quality (Fig. 6c). Next, we tested whether Docetaxel-TCO specifically binds to microtubules and enables live-cell labeling of the cytoskeleton with H-Tet-SiR. Live-cell standard confocal as well as super-resolution microscopy images recorded by re-scan confocal microscopy (RCM)^44,45^ and structured illumination microscopy (SIM)^46^ demonstrate specific labeling of the cytoskeleton by click chemistry (Fig. 6d). Furthermore, we performed live-cell re-scan confocal time lapse microscopy showing that microtubules exhibit damped dynamics lacking microtubule polymerization and degradation events as seen in our EMTB labeling approach when applying the microtubule-stabilizing antimitotic drug Docetaxel-TCO at micromolar concentrations (compare Supplementary Videos 1,2, and 3).

## Conclusions

With continuously increasing spatial resolution of super-resolution microscopy the development of small and efficient fluorescent probes generating minimal linkage errors becomes particularly important. Site-specific introduction of a ncAA such as TC0*-Lys into the amino acid chain of the protein of interest followed by bioorthogonal click chemistry with tetrazine-dyes represents a broadly useful possibility to overcome current limitations and enable high-end fluorescence imaging with organic dyes. In addition, a broad range of organic dyes are commercially available as tetrazine-dyes. Whereas FRET is responsible for fluorescence quenching in blue- and green-absorbing dyes, we discovered PET from the excited dye to tetrazine as the main quenching mechanism in red-absorbing oxazine and Si-rhodamine derivatives. We show that a stacked arrangement of the conjugated π-electron system of the dye and tetrazine is required for strong fluorescence quenching. Upon reaction with dienophiles quenching interactions are reduced resulting in a considerable increase in fluorescence intensity. Because tetrazine-dyes provide linkage error free stoichiometric labeling of proteins at well-defined extracellular and intracellular positions they are ideally suited for single-molecule fluorescence imaging and tracking as well as super-resolution microscopy experiments in fixed and living cells.

## Methods

### Tetrazine-dyes/TCO*-Lys

Me-Tet derivatives of ATTO425, ATTO465, ATTO488, ATTO550, ATTO565, ATTO590, ATTO594, ATTO620, ATTO655, ATTO680, ATTO700, AZ503, AZ519, and ATTO647N were provided by ATTO-TEC (Siegen, Germany). Me-Tet-ATTO532, H-Tet-Cy3, Me-Tet-TAMRA, H-Tet-Cy5 were used from Jena Bioscience (Jena, Germany). ATTO532, Cy3, TAMRA, Cy5. HMSiR-NHS was purchased from MoBiTec (Goettingen, Germany) and TCO*-Lys from SiChem (Bremen, Germany).

### DNA constructs and plasmids

Plasmid amplification was performed via transformation in E. coli (XL1-Blue) and DNA isolation via MIDI-prep (NucleoBond^®^ Xtra Midi, Macherey & Nagel, #740410). The respective amber stop mutants were generated by introducing a TAG codon through PCR-based site-directed mutagenesis. The plasmid for expression of the TNF-receptor pTNFR1-tdEOS was a gift from Mike Heilemann (Addgene plasmid # 98273). The plasmid for the expression of the Kainate receptor (pcDNA3-GluK2) was a gift from Dr. Peter Seeburg (Max-Planck Institute for Medical Research, Heidelberg, Germany). An AgeI restriction site was introduced at the C-terminus of the GluK2 coding sequence in front of the STOP-codon. Subsequently, GluK2 was cloned into the MCS of the pEGFP-N1 plasmid (Clontech #U55762) using the restriction enzymes EcoRI and AgeI. The plasmid for expression of the microtubule-associated protein Ensconsin (EMTB-3xGFP) was a gift from William Bement (Addgene plasmid # 26741). The plasmid for the expression of the tRNA/tRNA-synthetase pair (pCMV tRNA^Pyl^/NESPylRS^AF^) was kindly provided by Edward Lemke^54^.

#### Transfection

Cells were transfected at 60 - 80 % confluence with a commercial transfection reagent (JetPrime, Polypus, #114-01) with the suitable plasmid/reagent mixture (according to manufacturer´s recommendations). The total DNA amount per well was 500 ng in a 1:1 ratio between the DNA for the respective POI and pCMV-tRNAPyl/NESPylRSAF. The ncAA (TCO*A; SiChem GmbH, Bremen, #SC-8008) was supplemented separately at a final concentration of 250 μM, diluted 1:4 with 1 M HEPES. After 6 - 8 h the medium was exchanged to fresh cell growth medium. Cells were incubated approx. 12 - 36 h before labeling and fixation/imaging.

#### Bioorthogonal labelling

For bioorthogonal labeling of clickable receptors the transfected HEK293T or COS-7 cells were incubated with 1.5 μM of the respective tetrazine-dye in cell growth medium for 10 min at room temperature or on ice. After incubation the cells were washed twice and fixed at room temperature with a mixture of 4 % formaldehyde and 0.25 % glutaraldehyde for 10 minutes. The cells were again washed three times with Hank’s Balanced Salt solution (HBSS, Sigma, #55037C) before imaging. Alternatively, the cells were imaged live without fixation. For labelling of actin filaments via pre-labeling with phalloidin-TCO the cells were first fixed with 4% formaldehyde in cytoskeleton buffer (10 mM MES pH 6.1, 150 mM NaCl, 5 mM EGTA, 5 mM glucose, 5 mM MgCl2) at room temperature for 10 min. Cells were then washed 2-3 times with HBSS (ThermoFisher, #14025092) and then incubated for at least 1 h with phalloidin-TCO (~0.5 μM) at room temperature or alternatively for 8 h overnight at 4°C. Afterwards the cells were washed again 2-3 times with PBS and stored until further labeling/imaging. Fluorescence labeling was performed by adding the respective tetrazine-dye (1-3 μM in PBS) for 10 min at room temperature. Cells were then rinsed with 2-3 times PBS and imaged or, for wash-free applications, imaged without removing the tetrazine-dye solution.

#### *d*STORM

The *d*STORM images were acquired using an inverted wide-field fluorescence microscope (IX-71; Olympus). For excitation a 641nm diode laser (Cube 640-100C, Coherent, Cleanup 640/10, Chroma) was focused onto the back focal plane of the oil-immersion objective (60x, NA 1.45; Olympus). Emission light was separated from the illumination light using a dichroic mirror (635rpc, Chroma) and spectrally filtered by a bandpass filter (Em01-R442/514/647-25; Semrock). Images were recorded with an EMCCD (IXON DU897, Andor). Resulting pixelsize for data analysis was measured as 128 nm. For each *d*STORM measurement, at least 15,000 images with an exposure time of 20 ms and irradiation intensities of ~ 2 kW/cm^2^ were recorded by HILO (highly inclined and laminated optical sheet) illumination. Experiments were performed in PBS-based photoswitching buffer containing 100 mM β-mercaptoethylamine (MEA, Sigma-Aldrich) for Cy5 and an oxygen scavenger system (2 % (w/v) glucose, 4 U/ml glucose oxidase and 80 U/ml catalase) or only in PBS for Cy5B and HMSiR, adjusted to pH 7.4. Image reconstruction was performed using rapidSTORM3.3.

## Acknowledgements

We thank ATTO-TEC GmbH for the generous provision of Me-T et-ATTO-dyes. We thank Petra Gessner for providing expert technical assistance in immunocytochemistry, and cell culture preparation. We are grateful to Mike Heilemann for providing the TNFR plasmid. We thank Edward Lemke and Gemma Estrada Girona (European Molecular Biology Laboratory, Heidelberg, Germany) for the gift of the pCMV tRNA^Pyl^/NESPylRS^AF^ plasmid and expert training on how to use it. This work was supported by the Deutsche Forschungsgemeinschaft CRC-TRR 166 to M.S. (A04, B04), and S.D. (B02). EMTB-3XGFP was a gift from William Bement (Addgene plasmid # 26741).

## Author contributions

G.B. and M.S. conceived and designed the project. M.S. and S.D. supervised the project. G.B., A.K., L.B.-P., and A.K. performed all labeling and imaging experiments. Z.-D.S., M.Sch., J.B.G., and L.L., provided the dyes Cy5B and JF_646_, and corresponding tetrazine derivatives. N.W. and J.S. provided docetaxel-TCO. G.B., A.K. and S.D. performed data analysis. G.B., S.D., and M.S. wrote the manuscript.

## Additional information

Supplementary information and chemical compound information are available in the online version of the paper. Correspondence and requests for materials should be addressed to M.S.

## Competing financial interests

The authors declare no competing financial interests.

## Supplementary Information

### Chemical Synthesis of tet-HMSiR und tet-Cy5B and phalloidin-TCO

H-Tet-HMSiR0.034: μmol HMSiR-NHS (MoBiTec, Göttingen, Germany, # A208-01) was added to 0.68 μmol H-Tet-amine (Jena Bioscience, Jena, Germany, # CLK-001-5) in water-free DMSO (Thermo Fisher Scientific, Invitrogen Cat. Nr. D12345) and diisopropylethylamine (Sigma-Aldrich, # D125806, DIPEA, 1.1 μmol) and incubated for 4 h at room temperature. H-Tet-Cy5B: 0.355 μmol Cy5B-NHS^8^ was added to 3.5 μmol H-Tet-amine (Jena Bioscience, Jena, Germany, # CLK-001-5) in water-free DMSO (Thermo Fisher Scientific, Invitrogen Cat. Nr. D12345) with diisopropylethylamine (Sigma-Aldrich, # D125806, DIPEA, 1.1 μmol) and incubated for 4 h at room temperature. Phalloidin-TCO: 0.07μmol Amino-phalloidin (Biomol, Hamburg, Germany, #ABD-5302) was added to 0.7 μmol TCO-PEG4-NHS (Jena Bioscience, Jena, Germany, #CLK-A137-10) in water-free DMSO (Thermo Fisher Scientific, Invitrogen Cat. Nr. D12345) with diisopropylethylamine (DIPEA, 1.1 μmole) and incubated for 4 h at room temperature. Tetrazine-dyes and phalloidin-TCO were purified by reverse phase chromatography on a Phenomenex Kinetex Biphenyl Core-shell LC Columns, (2.6 μm, 150 x 4.6 mm; water:acetonitrile with 0.1% formic acid and an elution gradient of 0-95% in 45 min).

### Chemical synthesis of docetaxel-TCO probe

*General experimental information*. All chemical reagents for the synthesis were purchased from commercial suppliers and were used without further purification: Docetaxel (TCI, Eschborn, Germany, CAS: 114977-28-5); (E)-cyclooct-4-en *p*-nitrophenol active ester (SiChem, Bremen, Germany, CAS: 1438415-89-4); 6-aminohexanoic acid (Alfa Aesar, Karlsruhe, Germany, CAS: 60-32-2). Dimethylformamide and triethylamine were purchased from Sigma-Aldrich (Traufkirchen, Germany).

^1^H and ^13^C NMR spectra were recorded at 295 K on a Bruker Avance III HD 400 (^1^H: 400 MHz, ^13^C: 100 MHz) and Bruker Avance III HD 600 (^1^H: 600 MHz, ^13^C: 150 MHz). The chemical shifts (*δ*) are reported in parts per million (ppm) with respect to the solvent residual signals of CD3OD (*δ*(MeOD-d4) = 3.31 ppm for ^1^H and *δ*(MeOD-d4) = 49.00 for ^13^C). The coupling constants (J) are reported in Hz and indicate the proton spin-spin couplings. Multiplicity is abbreviated as follows: s = singlet; d = doublet; t = triplet; q = quartet; m = multiplet; dd = doublet of doublet; dt = doublet of triplet; br s = broad singlet, br d = broad doublet; br t = broad triplet etc.

High resolution mass spectrometry (HRMS) of synthesized compounds was confirmed with Bruker Daltonics micrOTOF-Q III with electrospray ionization (ESI).

Liquid chromatography purification was performed with silica gel 60 (0.04 - 0.063 mm), purchased from Macherey-Nagel (Düren, Germany).

*Synthetic procedure and characterization*. The synthesis of docetaxel-TCO (6) was performed in three steps (Scheme 1). Docetaxel (1) was deprotected with formic acid to obtain intermediate 2 as previously reported.^[1]^ The reaction of 6-aminohexanoic acid (3) and (*E*)-cyclooct-4-en *p*-nitrophenol active ester (4) in DMF/H_2_O mixture overnight led to (*E*)-cyclooct-4-en-1-yl-*N*-hexanoic acid carbamate (5). The subsequent amide coupling between intermediates 2 and 5 generated the docetaxel-TCO probe 6.

**Supplementary Scheme 1:**
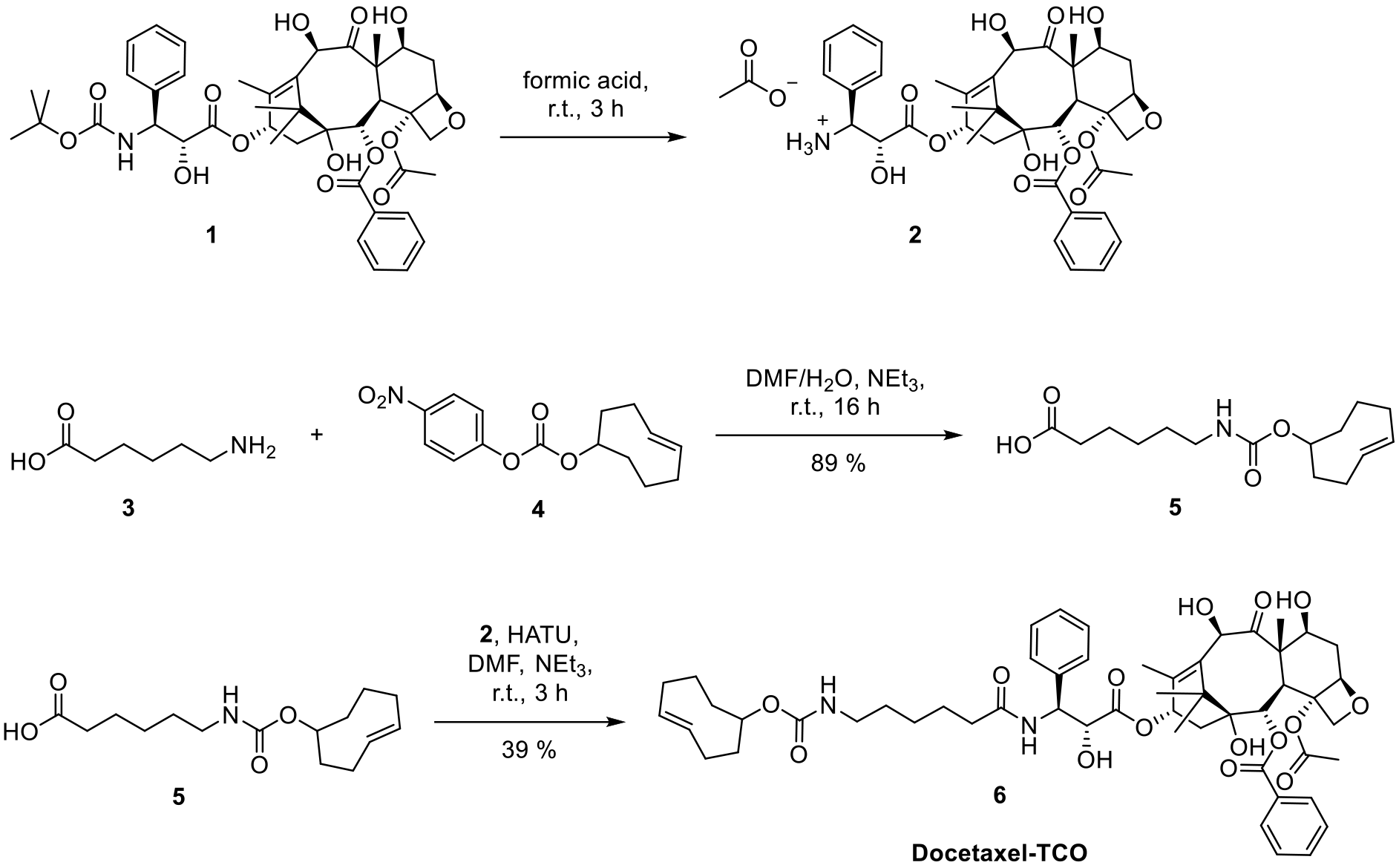
Synthesis of docetaxel-TCO probe.

*3’-Aminodocetaxel (2)*

In a 5 mL flask docetaxel (1) (100 mg, 124 μmol, 1 eq.) was dissolved in 0.5 mL formic acid and stirred at r.t. for 3 h. Afterwards the solvent was evaporated to dryness under reduced pressure and the obtained crude formate salt 2 was used in the next step without further purification.

*(E)-Cyclooct-4-en-1-yl-N-hexanoic acid carbamate (5)*

In a 5 mL flask 6-aminohexanoic acid (3) (25.0 mg, 191 μmol, 1 eq.) was suspended in 0.5 mL DMF and 0.1 mL H2O and 132 μL NEt_3_ (953 μmol, 5 eq.) were added. After stirring for 5 min (E)-cyclooct-4-en *p*-nitrophenol active ester (4) (55.5 mg, 191 μmol, 1 eq.) was added and the reaction mixture was stirred overnight. The clear yellow solution was diluted with 50 mL AcOEt and 10 mL 1 m NH4Cl. The aqueous phase was separated and the organic phase was washed once with 10 mL brine. The organic layer was dried over Na2SO4 and concentrated under reduced pressure. The residue was purified by column chromatography using a gradient: CHCl3 → CHCl3:MeOH = 25:1. The TCO-Carbamate 5 was obtained as colorless oil (48 mg, 89%).

**^1^H NMR** (400 MHz, MeOD-*d_4_*): *δ* = 5.72-5.64 (m, 1H), 5.59-5.50 (m, 1H), 4.83-4.79 (m, 1H), 3.10 (t, *J* = 6.7 Hz, 2H), 2.42-2.21 (m, 6H), 1.89-1.23 (m, 12H).

**^13^C NMR** (100 MHz, MeOD-*d_4_*): *δ*= 177.7, 158.9, 136.5, 132.7, 71.4, 42.1, 41.7, 35.4, 35.0, 33.9, 31.0, 30.8, 29.1, 27.6, 25.9.

**HRMS** (ESI): *m/z* calc. for C_15_H_25_NNaO_4_^+^ [M+Na]^+^: 306.1676; found: 306.1683 (Δ = 2.4 ppm). *Docetaxel-TCO probe (6)*

To a solution of TCO-carbamate 5 (30.0 mg, 106 μmol, 1 eq.) in 0.5 mL dry DMF successively HATU (44.3 mg, 117 μmol, 1.1 eq.) and NEt_3_ (74 μL, 5 eq.) were added. After stirring for 10 min at r.t. 3’-aminodocetaxel formate salt 2 (93.5 mg, 122 μmol, 1.15 eq.) was added and the reaction mixture was further stirred for 3 h. After full consumption of TCO-carbamate 5 the reaction mixture was diluted with 50 mL AcOEt and 10 mL 1 m NH4Cl. The aqueous phase was separated and the organic phase was washed once with 10 mL brine. The organic layer was dried over Na2SO4 and concentrated under reduced pressure. The residue was purified by column chromatography twice using a gradient: (CHCl_3_:MeOH = 25:0.25 → CHCl_3_:MeOH = 25:1). The docetaxel-TCO probe (6) was obtained as colorless solid (40.2 mg, 39 %).

**^1^H NMR** (400 MHz, MeOD-*d*4): *δ*= 8.12-8.11 (m, 1H), 8.11-8.10 (m, 1H), 7.68-7.64 (m, 1H), 7.58-7.54 (m, 2H), 7.44-7.37 (m, 4H), 7.30-7.25 (m, 1H), 6.16 (t, *J* = 9.1 Hz, 1H), 5.70-5.45 (m, 4H), 5.27 (s, 1H), 4.99 (dd, *J* = 9.0, 0.7 Hz, 1H), 4.81-4.77 (m, 1H), 4.58 (d, *J* = 4.6 Hz, 1H), 4.24-4.17 (m, 3H), 3.87 (d, *J* = 7.2), 3.05 (br t, *J* = 6.9, 2H), 2.44 (ddd, *J* = 14.4, 9.7, 6.5 Hz, 1H), 2.39-2.14 (m, 10H), 2.10-1.98 (m, 2H), 1.90 (d, *J* = 1.2 Hz, 3H), 1.87-1.80 (m, 2H), 1.76-1.59 (m,7H), 1.57-1.28 (m, 6H), 1.19 (s, 3H), 1.13 (s, 3H).

**^13^C NMR** (100 MHz, MeOD-*d*4): *δ*= 211.1, 175.9, 174.4, 171.8, 167.7, 158.7, 158.6, 140.1, 139.2, 138.0, 136.3, 134.5, 132.5, 131.4, 131.2, 129.7, 129.7, 128.9, 128.4, 85.9, 82.3, 79.1, 77.6, 76.4, 75.6, 74.8, 72.7, 72.5, 71.2, 58.9, 56.8, 47.8, 44.5, 42.0, 41.5, 37.5, 36.9, 36.8, 35.3, 35.2, 33.7, 30.9, 30.7, 29.0, 27.4, 27.1, 26.7, 26.6, 26.0, 23.2, 21.7, 14.4, 10.5.

**HRMS** (ESI): *m/z* calc. for C_53_H_69_N_2_O_15_^+^ [M+H]^+^: 973.46925; found: 973.46794 (Δ = 1.34 ppm).

### Measurements of absorbance/emission spectra

Time-dependent fluorescence intensities were measured in quartz glass cuvettes using a FP-6500 spectrofluorimeter (Jasco). Fluorescence was excited at the absorption maxima. The sample temperature was adjusted to 25°C using a Peltier thermocouple.

### Time-correlated single-photon counting (TCSPC)

All fluorescence lifetime measurements at 640 nm excitation were performed with a FluoTime200 time resolved spectrometer (Picoquant, Berlin, Germany). Decay curves were analyzed using FluoFit 4.4.0.1 software. Measurements of fluorescence lifetimes of dyes with excitation at 400 - 560 nm were carried out with a MicroTime200 (Picoquant, Berlin, Germany), equipped with a supercontinuum laser SuperK Extreme EXW12 (NKT Photonics, Germany). The setup is attached to an Olympus IX83 with a 60x/1.2 ultra-plan-apochromat water-immersion objective.

### Intermolecular fluorescence quenching experiments

To interpret quenching efficiencies of tetrazine and fluorophores, bimolecular quenching experiments with ATTO655, ATTO700, JF_646_, and Cy5 were performed with Me-Tet-amine in phosphate-buffered saline (PBS) at pH 7.4. Bimolecular static and dynamic quenching constants, *K_stat_* and *K_dyn_*, were determined from time-resolved and steady-state fluorescence quenching experiments using Stern-Volmer analysis:

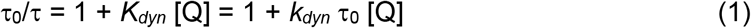

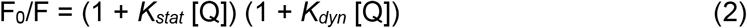

where τ_0_ and F0 are the fluorescence lifetime and intensity in the absence of a quencher, τ and F are the fluorescence lifetime and intensity in the presence of the quencher Q with the concentration [Q], and *K_stat_* and *K_dyn_* denote the static and dynamic Stern-Volmer constant, respectively. Dynamic quenching is attributed to collisional interactions at a dynamic quenching rate constant of *k_dyn_·* Static quenching originates in bimolecular complexes that are very efficiently quenched and *K_stat_* can thus be interpreted as an association constant *K_a_*. Experimental estimates are provided as results from linear regression together with standard error from the fitting procedure.

### Fluorescence-Correlation-Microscopy (FCS)

PET-FCS was performed using a custom-built confocal fluorescence microscope setup as described previously^39,40^ equipped with a 640 nm diode laser (OBIS640, Coherent Europe B.V., Utrecht, Netherlands). Samples were diluted in PBS at pH 7.4 and filtered through a 0.2 μm syringe filter, transferred onto a microscope slide and covered by a cover slip. Samples with concentrations of 1 nM yielded an average of ~20 molecules in the detection focus of the microscope setup. Sample temperature was adjusted to 25 °C using a custom-built objective heater. For each sample, at least three individual autocorrelation functions were recorded of at least 5 min measurement time each. PET-FCS data was fitted to an analytical model for diffusion and a two-state equilibrium with off- and on-rate constants *k_closing_* and *k_opening_*:

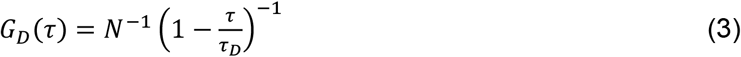

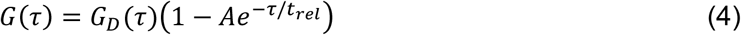

where *N* is the number of detected molecules; *τd* is the diffusion time; *A* = *k_closing_ / k_opening_* is the equilibrium constant and *τ_rel_* = (*k_closing_* + *k_opening_*)^-1^ is the exponential time constant.

### Confocal microscopy

Confocal fluorescence images were recorded on a commercial LSM700 (Zeiss; 63x/1,4 oil objective). Single plane images were acquired with a pixel size of 90nm and suitable settings for the respective dye. Images were adjusted for brightness and contrast (only linear changes).

### Rescan confocal microscopy (RCM)

RCM imaging was performed using the commercially available RCM-unit (confocal.nl) attached to an inverse Nikon TiE microscope body. The system is equipped with a multiline (405 nm, 488 nm, 561 nm, 640 nm) laser module Skyra (Cobolt) and a sCMOS camera Zyla4.2P (Andor). Images were acquired with an 100x/1.49 oil objective and a resulting pixel size of 42 nm. Images were corrected for the camera offset and a median filter (kernel: 1px) was applied.

### Structured illumination microscopy (SIM)

SIM imaging was performed on a commercial ELYRA S.1 microscope (Zeiss AG). The setup is equipped with a Plan-Apochromat 63x/1.40 immersion-oil based objective and four excitation lasers, a 405 nm diode (50 mW), a 488 nm OPSL (100 mW), a 561 nm OPSL (100 mW) and a 642 nm diode laser (150 mW).

### Cell culture and sample preparation

HEK293T cells (German Collection of Microorganisms and Cell Cultures, Braunschweig, Germany; #ACC635) were maintained in T25-culture flasks (Thermo Fisher Scientific, Cat. Nr. 156340) with Dulbeccos´s Modified Eagle´s Medium (DMEM, Sigma Aldrich, #D5796) with 10 % FCS (Sigma-Aldrich, #F7524), 2 mM L-Glutamine (already in DMEM suppl.) and 100 U/mL Penicillin + 0.1 mg/ml Streptomycin (Sigma-Aldrich, #P4333) in a 5 % CO_2_ atmosphere at 37 °C.

COS-7 cells (Cell Lines Service GmbH, Eppelheim, Germany #605470) were maintained in DMEM (Sigma, #D8062) with 10% FCS (Sigma-Aldrich, #F7524), 2 mM L-Glutamine (already in DMEM suppl.) and 100 U/ml Penicillin + 0.1 mg/mL Streptomycin (Sigma-Aldrich, #P4333) in a 5% CO_2_ atmosphere at 37°C. NG108-15 cells (ATCC HB-12317) were maintained in DMEM (Sigma, #D8062) with 10% FCS (Sigma-Aldrich, #F7524), 2mM L-Glutamine (already in DMEM suppl.) and 100U/mL Penicillin+0,1mg/mL Streptomycin (Sigma-Aldrich, #P4333) in a 5% CO_2_ atmosphere at 37°C.

NIH-3T3 (German Collection of Microorganisms and Cell Cultures, Braunschweig, Germany; #ACC59) were maintained in DMEM (Sigma, #D8062) with 10% FCS (Sigma-Aldrich, #F7524), 2 mM L-Glutamine (already in DMEM suppl.) and 100 U/ml Penicillin + 0.1 mg/ml Streptomycin (Sigma-Aldrich, #P4333) in a 5% CO_2_ atmosphere at 37°C. NG108-15 cells (ATCC HB-12317) were maintained in DMEM (Sigma, #D8062) with 10% FCS (Sigma-Aldrich, #F7524), 2mM L-Glutamine (already in DMEM suppl.) and 100 U/ml Penicillin + 0.1 mg/ml Streptomycin (Sigma-Aldrich, #P4333) in a 5% CO_2_ atmosphere at 37°C.

U2-OS cells (Cell Lines Service GmbH, Eppelheim, Germany # 300364) were maintained in DMEM (Sigma, #D8062) with 10% FCS (Sigma-Aldrich, #F7524), 2 mM L-Glutamine (already in DMEM suppl.) and 100 U/ml Penicillin + 0.1 mg/ml Streptomycin (Sigma-Aldrich, #P4333) in a 5% CO_2_ atmosphere at 37°C. NG108-15 cells (ATCC HB-12317) were maintained in DMEM (Sigma, #D8062) with 10% FCS (Sigma-Aldrich, #F7524), 2mM L-Glutamine (already in DMEM suppl.) and 100 U/ml Penicillin + 0.1 mg/ml Streptomycin (Sigma-Aldrich, #P4333) in a 5% CO_2_ atmosphere at 37°C.

Prior to seeding the cells, dishes were coated with 0.01% poly-D-Lysine (Sigma-Aldrich, #P6407) for 1 h at room temperature. For all live and fixed cell images, cells were seeded at least 16 h before transfection on 4-well Lab-Tek II chambered cover glasses (Nunc, cat. no. 155409).

---

**Supplementary Figure S1.**
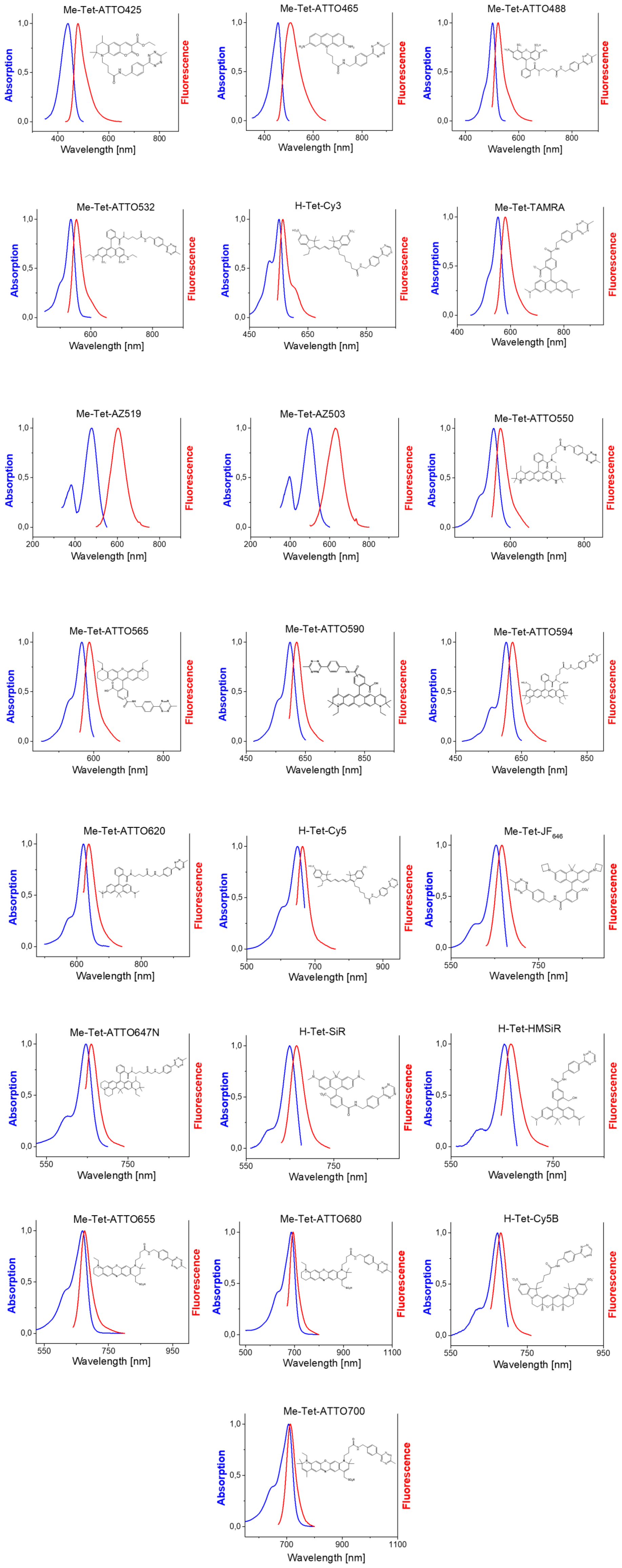
Molecular structure (according to availability) and absorption and emission spectra of all 22 tetrazine-dyes investigated in the study.

**Supplementary Figure S2.**
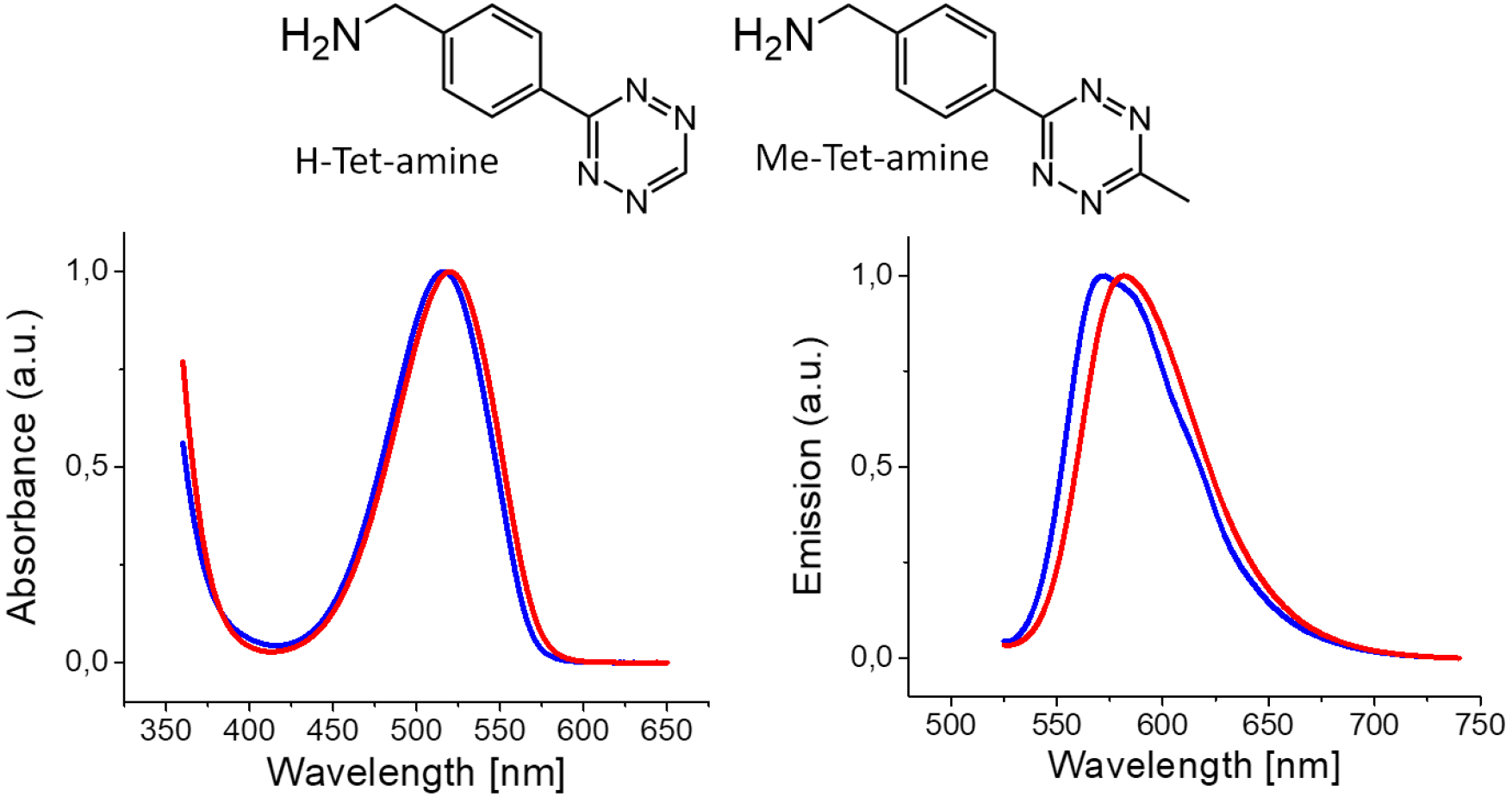
Molecular structure and absorption and emission spectra of H-Tet-amine (blue) and Me-Tet-amine (red).

**Supplementary Figure S3.**
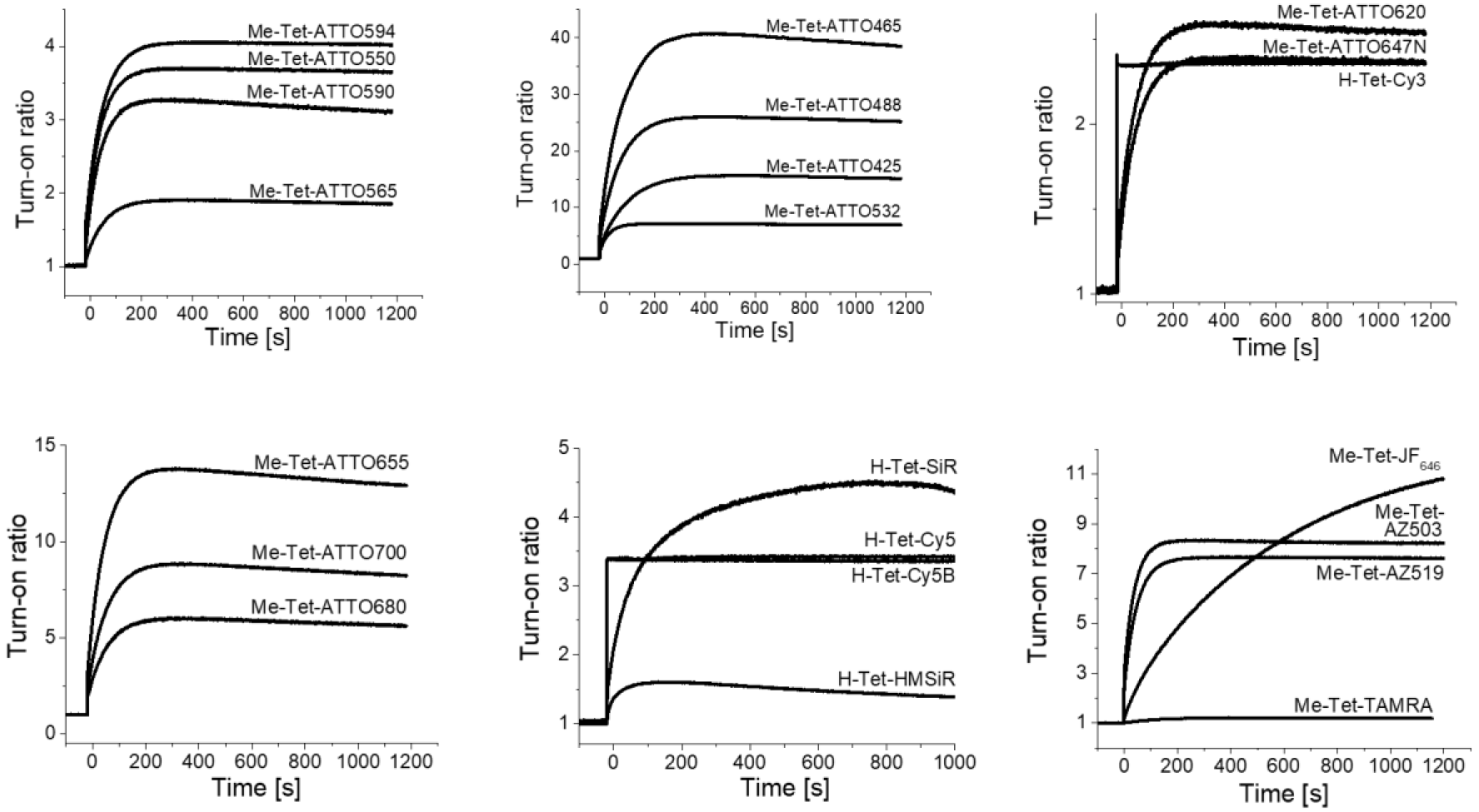
Fluorescence turn-on of 1 μm solutions of different tetrazine-dyes upon addition of 25 μM TCO*-Lys.

**Supplementary Figure S4.**
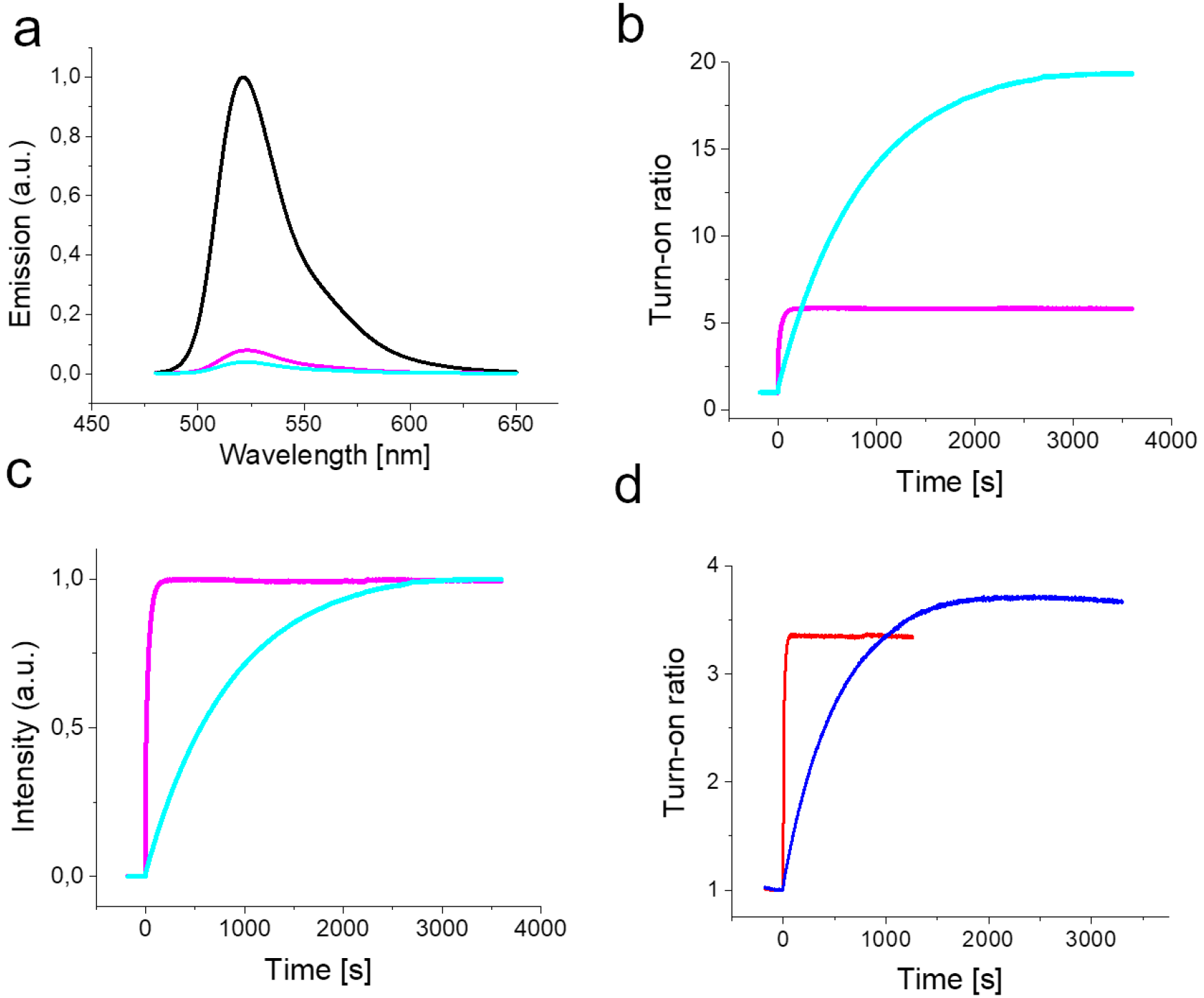
**a**. Relative fluorescence emission spectra of ATTO488 (black), H-Tet-ATTO488 (magenta), and Me-Tet-ATTO488 (light blue). The emission spectra are normalized to the same extinction at the excitation wavelength. **b**. Relative fluorescence intensity increase of H-Tet-ATTO488 (magenta) and Me-Tet-ATTO488 (light blue) upon addition of a 25-fold excess of TCO*-Lys measured at 522 nm and normalized to the fluorescence intensity recorded before addition of TCO*-lysine. **c**. Relative fluorescence intensity increase of H-Tet-ATTO488 (magenta) and Me-Tet-ATTO488 (light blue) upon addition of a 25-fold excess of TCO*-Lys normalized to the fluorescence intensity recorded after reaction with TCO*-Lys. **d**. Relative fluorescence intensity increase of H-Tet-Cy5 (red) and Me-Tet-Cy5 (blue) upon addition of a 25-fold excess of TCO*-lysine normalized to the fluorescence intensity recorded before addition of TCO*-Lys. All measurements were performed in aqueous buffer, PBS, pH 7.4.

**Supplementary Figure S5.**
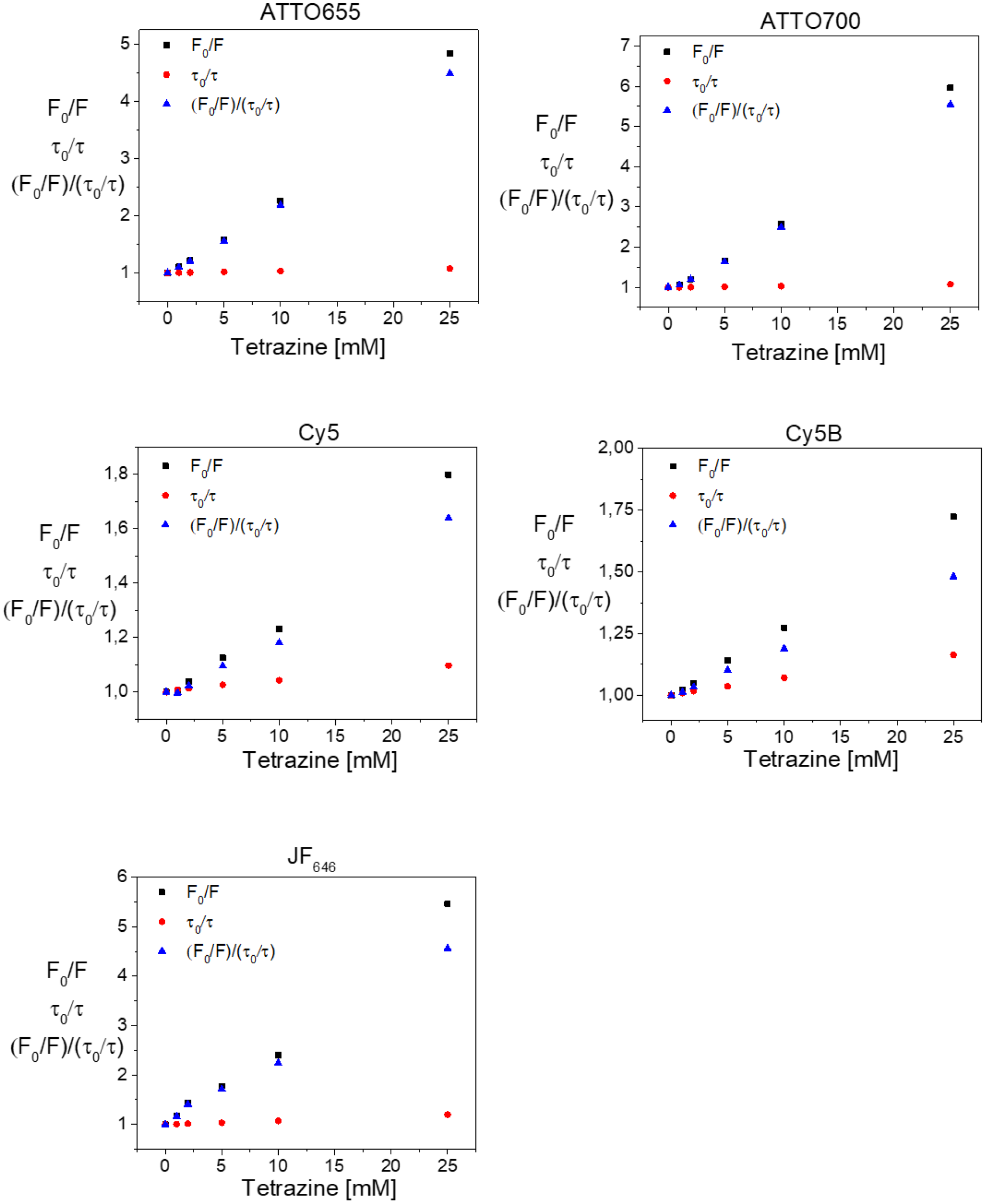
Steady-state and time-resolved Stern-Volmer plots of ATTO655, ATTO700, ATTO646, Cy5, Cy5B, and JF_646_ recorded with different concentrations of Me-Tet-amine (0 - 25 mM) in PBS, pH 7.4.

**Supplementary Figure S6.**
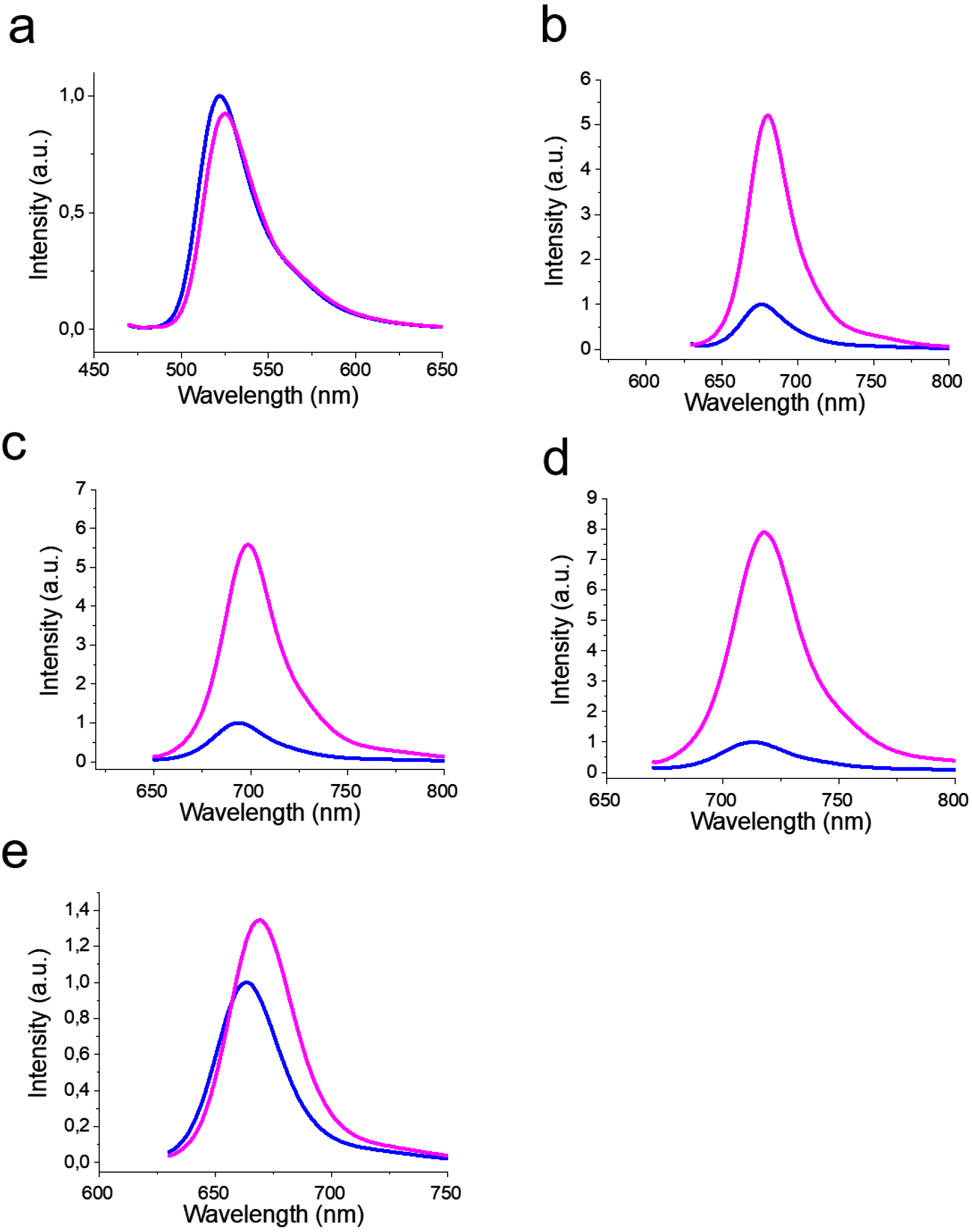
Relative fluorescence emission spectra of **a**. Met-Tet-ATTO488, **b**. Met-Tet-ATTO655, **c**. Met-Tet-ATTO680, **d**. Met-Tet-ATTO700, and **e**. H-Tet-Cy5 in PBS (pH 7.4) without (blue) and with 6M guanidiniumchlorid (magenta).

**Supplementary Figure S7.**
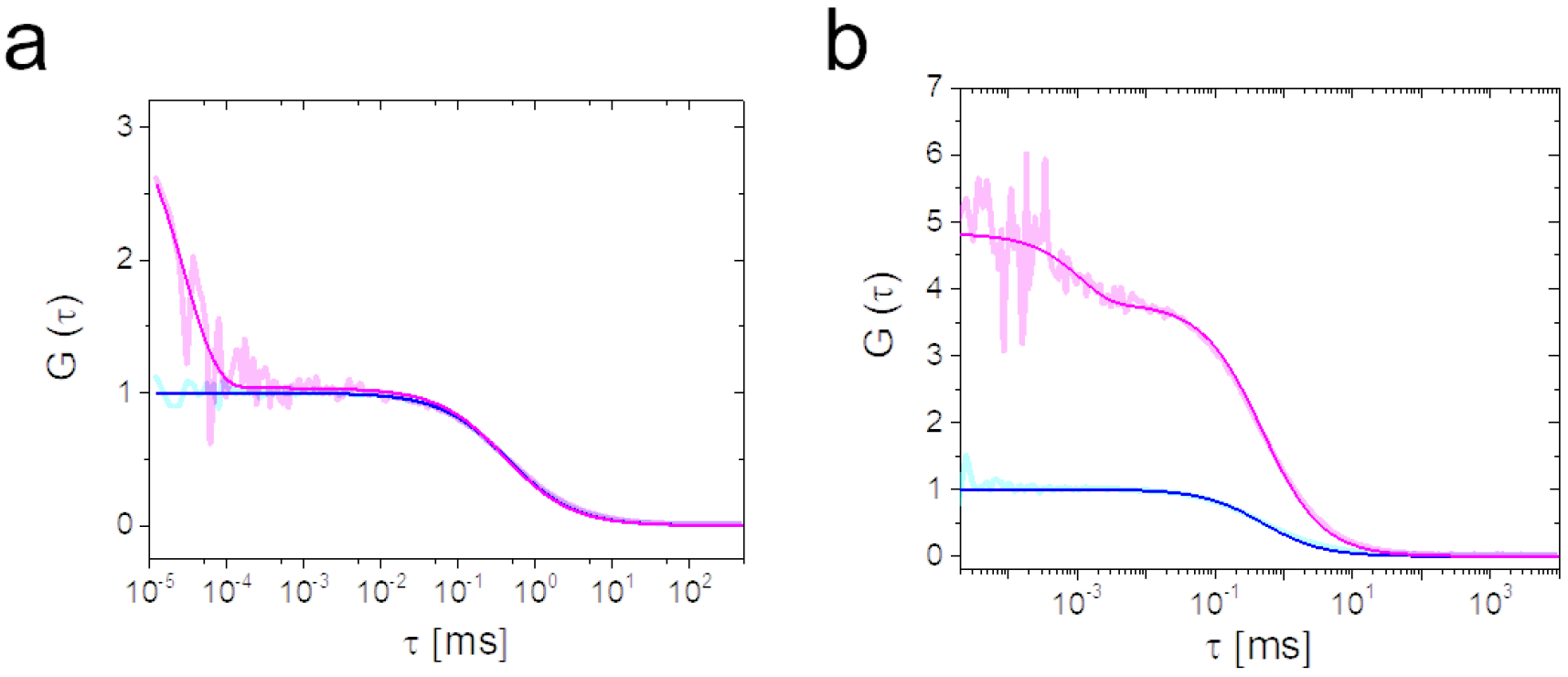
**a**. FCS curves of free ATTO655 alone (cyan) and in the presence of 25 mM Me-Tet-amine (magenta). **b**. FCS curves of Me-Tet-ATTO655 (magenta) and free ATTO655 (cyan). All data were recorded with dye concentrations of 1 nM and the FCS curves and normalized to the amplitude of the free ATTO655 data. Amplitude differences thus reflect the existence of quenched complexes that are stable on time scales longer than the diffusion time scale.

**Supplementary Figure S8.**
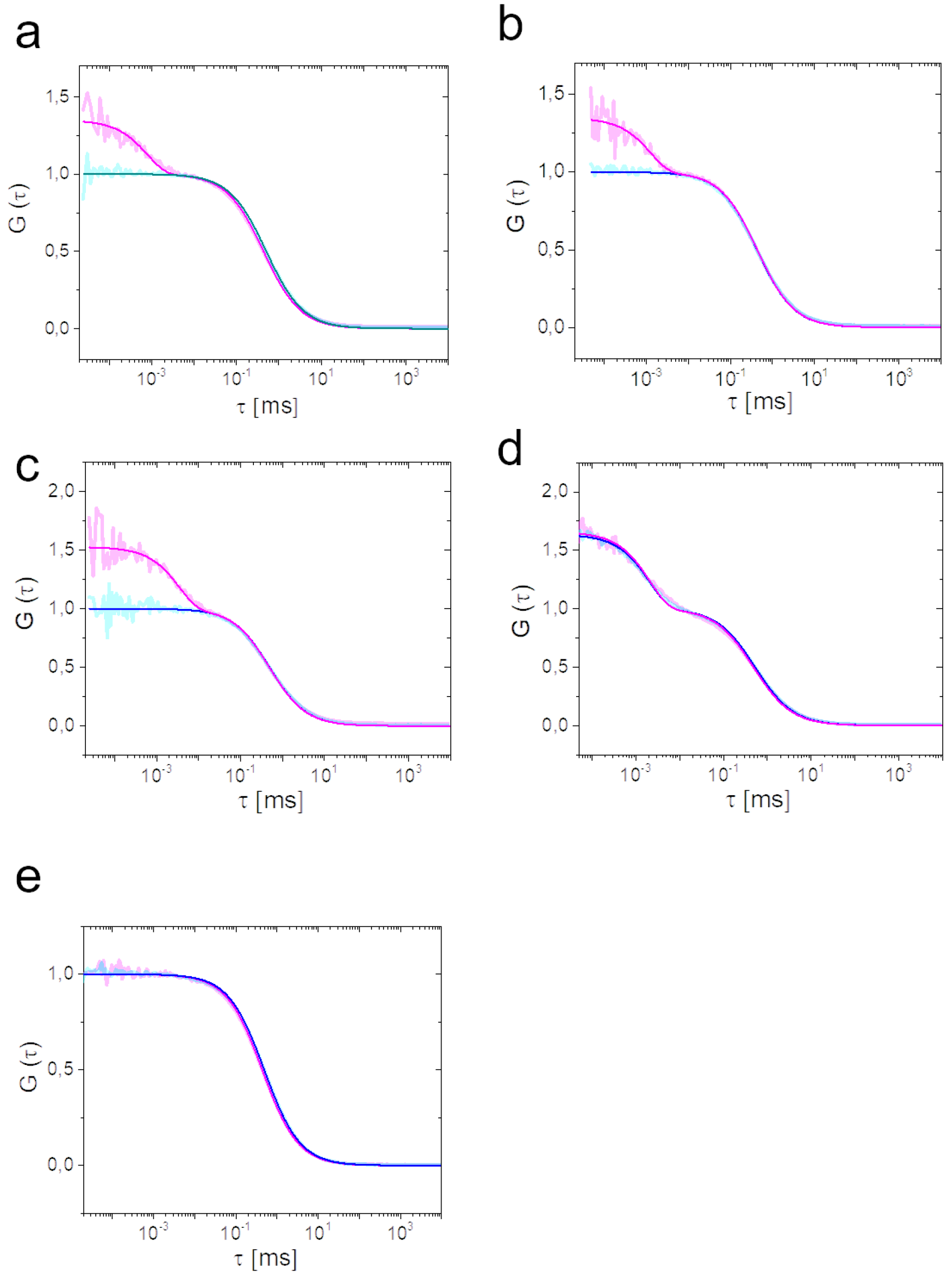
FCS curves of different tetrazine-dyes before (blue) and after clicking to TCO*-Lys (magenta). **a**. Met-Tet-ATTO655, **b**. Met-Tet-ATTO680, **c**. Met-Tet-ATTO700, **d**. H-Tet-Cy5, and **e**. H-Tet-Cy5B.

**Supplementary Figure S9.**
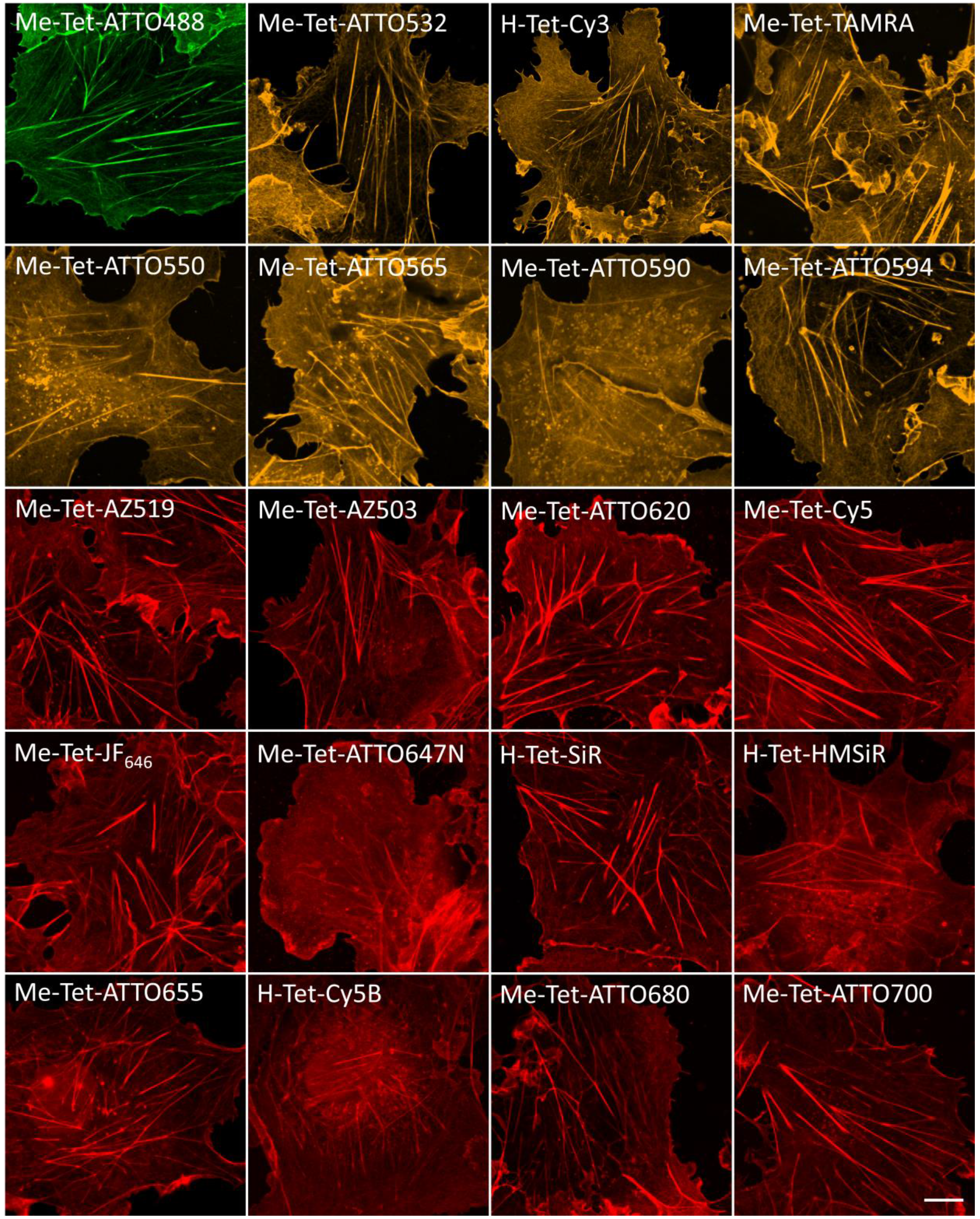
Confocal fluorescence images of COS7 cells labeled with phalloidin-TCO and different tetrazine-dyes. Cells were fixed and pre-labeled with an excess of phalloidin-TCO. Labeling was performed after washing with 1-3 μM tetrazine-dyes for 10 min. Before imaging, the excess of tetrazine-dyes was removed my washing with PBS, pH 7.4.

**Supplementary Figure S10.**
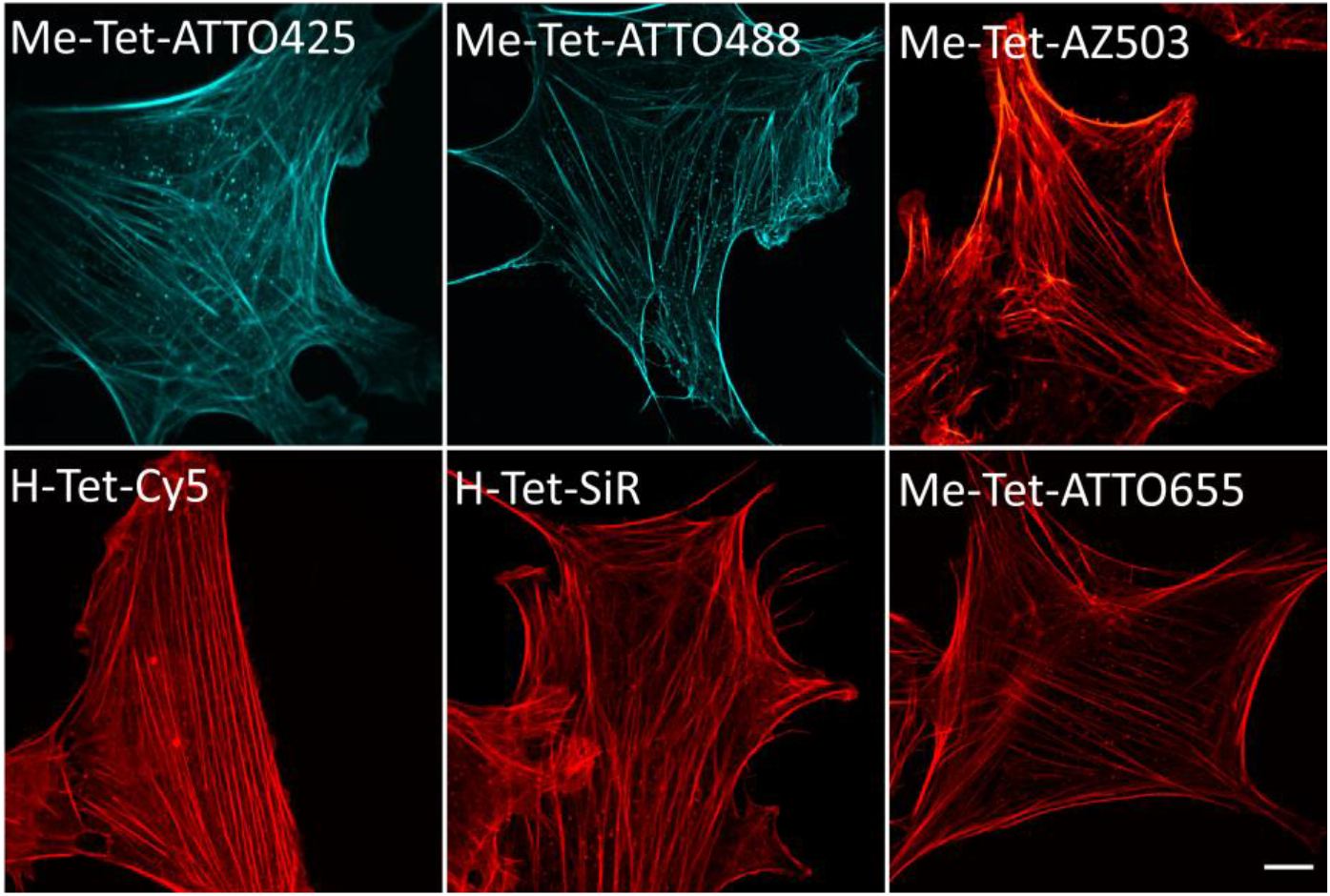
Re-scan confocal fluorescence images of NIH/33T3 cells labeled with phalloidin-TCO and different tetrazine-dyes. Cells were fixed and pre-labeled with an excess of phalloidin-TCO. Labeling was performed after washing with 1-3 μM tetrazine-dyes for 10 min. Before imaging, the excess of tetrazine-dyes was removed my washing with PBS, pH 7.4.

**Supplementary Figure S11.**
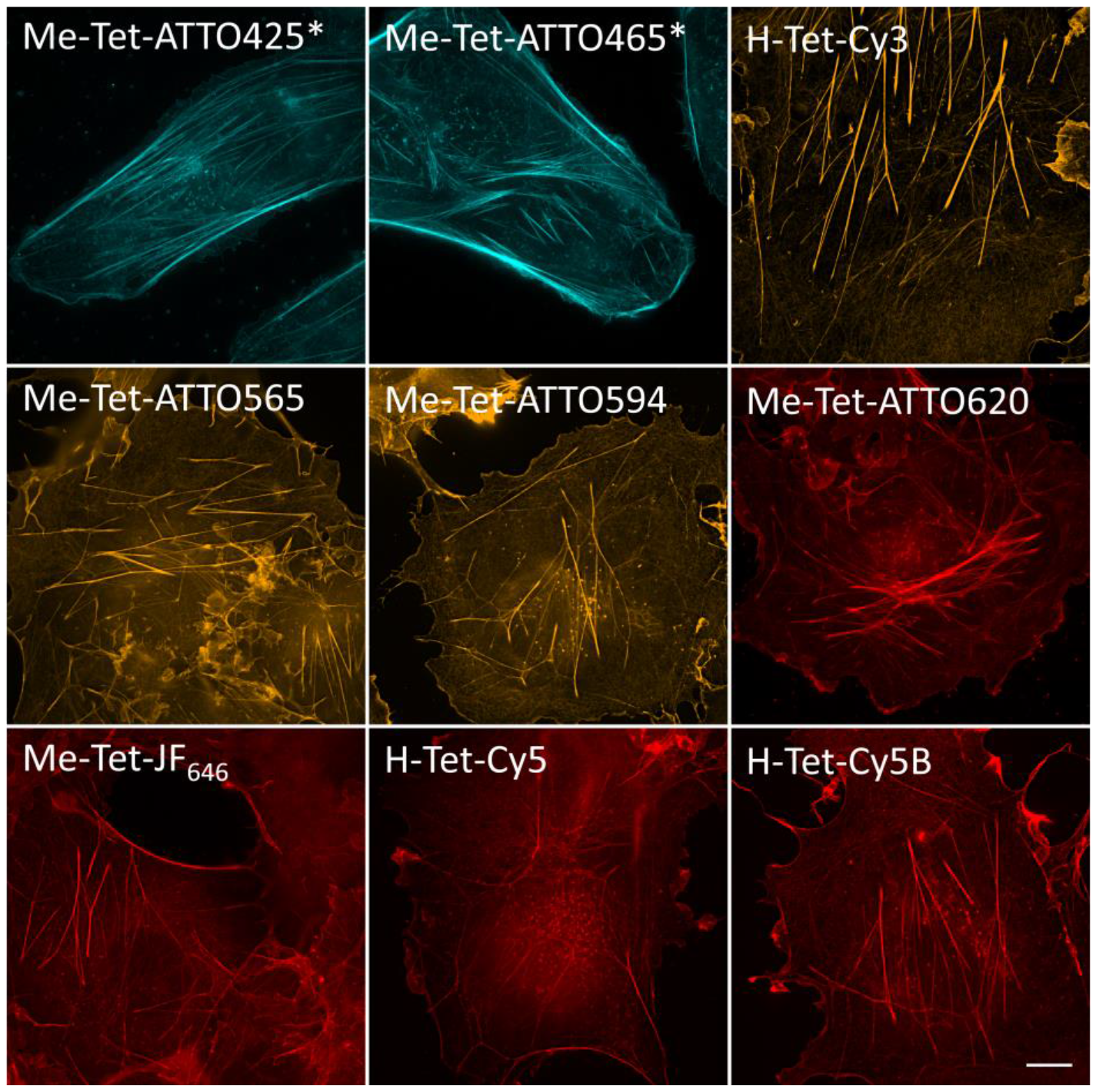
SIM images of U2-OS* and COS7 cells labelled with phalloidin-TCO and different tetrazine-dyes. Cells were fixed and pre-labeled with an excess of phalloidin-TCO. Labeling was performed after washing with 1-3 μM tetrazine-dyes for 10 min. Before imaging, the excess of tetrazine-dyes was removed my washing with PBS, pH 7.4.

**Supplementary Figure S12.**
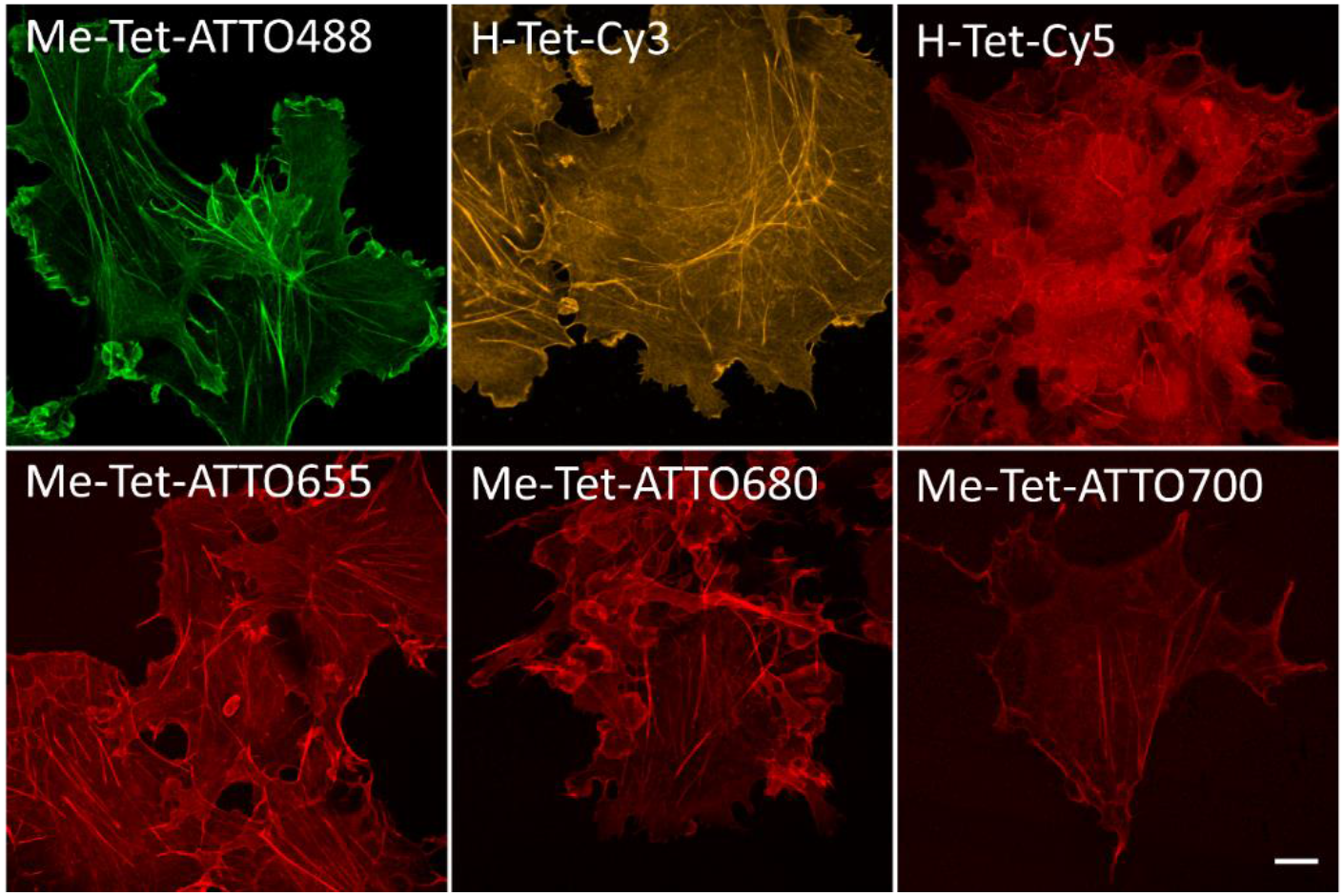
Wash-free confocal fluorescence images of COS7 cells labeled with phalloidin-TCO and different tetrazine-dyes. Cells were fixed and pre-labeled with an excess of phalloidin-TCO. Labeling was performed after washing with 1-3 μM tetrazine-dyes for 10 min in PBS, pH 7.4. without any additional washing step.

**Supplementary Videos 1 and 2.** Live-cell re-scan confocal time lapse microscopy of COS7 cells transfected with EMTB87TAG-3xGFP and clicked with 3 μM H-Tet-SiR for 10 min (GFP: green, SiR: magenta, and overlay). Cells were imaged in fresh cell culture medium.

**Supplementary Video 3.** Live-cell re-scan confocal time lapse microscopy of U2OS cell treated with 10 μM Docetaxol-TCO for 30 min and labelled with 10 μM H-Tet-SiR for 10 min. Cells were imaged in fresh cell culture medium.

Scale bars in all videos: 5 μm

Video speed: 20x

